# How well is Italian biodiversity represented in red lists and conservation legislation? Taxonomic biases, coverage gaps and the assessment-to-legislation bottleneck

**DOI:** 10.64898/2026.06.30.730850

**Authors:** Emanuele Miccolis, Maria B. Rasotto, Fabio De Pascale, Telmo Pievani

## Abstract

Italy, one of Europe’s most biodiverse countries, is expected to undergo landscape-wide nature recovery thanks to the EU Nature Restoration Regulation. National red lists represent critical instruments for informing conservation strategies at finer geographic resolutions and prioritizing taxa of national importance; yet the taxonomic completeness of Italian red lists remains unquantified in relation to national biodiversity. Equally, no systematic evaluation has been conducted on the representation of threatened and endemic Italian taxa within European and international conservation directives and treaties.

We present the first comprehensive review of threat status and policy inclusion of Italian biodiversity, encompassing animals, plants, fungi, lichens, and algae. We cross-referenced national species checklists with Italian, European, Mediterranean, and global IUCN red lists, alongside policy annexes from the Birds and Habitats Directives, the Bern and Barcelona Conventions, and CITES. Our dataset comprised 76,845 taxa, of which 8,389 are endemic; yet red list assessments exist for only 10% of this total (7,349 taxa, including 1,700 endemics).

Conservation policy coverage is even more restrictive: only 1,346 taxa are listed under at least one legislative instrument, with only half of these classified as threatened. This constitutes a compounding double bottleneck with most of the Italian biodiversity remaining both unassessed and unprotected and systematically biasing conservation policy toward an unrepresentative fraction of national biodiversity. We recommend accelerated national assessments and urgent establishment of a national biodiversity priority list founded on transparent prioritization protocols. This would complement ecosystem-based interventions mandated by the Nature Restoration Regulation while correcting for vertebrate-centred biases.

## 1. Introduction

Biodiversity worldwide is undergoing accelerated human-led extinctions and endangerment, a phenomenon thought to announce a sixth mass extinction event. This process of biodiversity impoverishment is exacerbated by a progressive loss of genetic diversity and complexity of species interactions in ecosystems (Ceballos et al., 2015; Cowie et al., 2022; Shaw et al., 2025). In June 2024, the European Union approved the legally-binding Nature Restoration Regulation (NRR), according to which every European member country is called to restore the 20% of terrestrial and marine natural areas by 2030, and overall builds upon existing legislations like the Habitats and Birds Directive (European Commission, 2024). The NRR complements the EU’s Biodiversity Strategy, aligning European restoration targets with the international commitments of the Kunming-Montreal Global Biodiversity Framework (CBD, 2022). One of the priorities is reducing the risk of extinction of species and preserving their adaptive potential, with the International Union for Conservation of Nature (hereafter IUCN) setting global rigour for the assessment of threat status. In addition, the IUCN supervises continent-wide and national red lists, which represent a more indicative tool for country-level prioritization based on risk of extinction.

National red lists are a valuable advancement for the assessment of the extinction risk of species at less coarse geographical scales, and local committees can opt for more specific assessments of endemic taxa and subspecies as they can uncover significant local declines of populations differently from global assessments. On the other hand, national red lists may not reflect global or pan-regional assessments due to differences in organizational structures and the inconsistent application of evaluation criteria. (Bender et al., 2012; Brito et al., 2010; Hilton-Taylor et al., 2000; Milano et al., 2021; Moon et al., 2023).

Inconsistencies in assessments across red lists may occur due to taxonomic impediments (i.e. the lack of taxonomic expertise and infrastructure to describe new taxa at the pace of their potential extinction) and taxonomic bias, defined even provocatively in the past as “taxonomic chauvinism” (Bonnet et al., 2002), which refers to the prioritization in conservation of charismatic taxa with limited connection to their role in the ecosystem or to their evolutionary uniqueness and threat status (Donaldson et al., 2017; Titley et al., 2017; dos Santos et al., 2020). Moreover, arbitrary choices in assessments may be transposed once taxa are added to conservation policies. The Habitats Directive presents prominent biases towards vertebrates and the preferred insects for inclusion are often large-bodied taxa with aesthetic value (Cardoso, 2012; Moser et al., 2016; Leandro et al., 2017; Tang and Visconti, 2021).

In Italy, the goals of international environmental frameworks are reprised in the National Strategy for Biodiversity 2030 (MASE, 2025), where national red lists are used as reference for objectives focused on stabilizing populations of threatened species.

Italy is one of the countries in the European Union with the highest richness of biodiversity according to the Convention on Biological Diversity (CBD, 2022), and new endemic species are constantly being described. While Italy invests in the conservation of endemic taxa via EU-funded projects (e.g. the EndemixIT project aiming to protect genomic heritage and integrity of five Italian endemics; https://endemixit.com/), there are currently no priority lists of endemic species. Red lists are not a standalone tool for prioritization in conservation planning; however, they are a main criterion in publishing priority lists. In priority scenarios, endemic status is amongst the highest ranks of prioritization as countries hold unique responsibility to avoid global extinction even when IUCN status has not been calculated yet (Fitzpatrick et al., 2007; Schmeller et al., 2008b, 2014; Kraus et al., 2023).

Italy was also the country that has published the first checklist of its fauna in Europe (Minelli and Stoch, 2007, https://www.faunaitalia.it/checklist/), and IUCN Italy has released several national red lists (https://www.iucn.it/). While these lists attempt to mention “Data Deficient” or “Not Evaluated” categories, the full extent of exclusion compared to national checklists is yet to be clarified. Furthermore, the proportion of Italian taxa currently integrated into international policy frameworks has yet to be systematically evaluated. Global (Bouchet et al., 1999; Challender et al., 2023), European (Moser et al., 2016), Mediterranean (Verlaque et al., 2019) and national reviews (Moreno-Saiz et al., 2021; Rossi et al., 2016) have highlighted how policies can misalign from red lists, either by attributing more optimistic evaluation or by omitting threatened taxa.

Considering the pivotal importance of Italy in protecting the biodiversity of the Mediterranean biodiversity hotspot, (Medail and Quezel, 1997; Médail and Quezél, 1999; Myers et al., 2000; Hewitt, 2011), in the present study we aim to provide a comprehensive account of assessed and non-assessed taxa across Italian biodiversity and IUCN lists in Europe. We subsequently mapped the legal protection of each taxon across major international frameworks. This multi-scale approach helped identify gaps in red lists, check if international assessments offset deficiencies in Italian assessments or, conversely, where Italian lists succeed in ensuring detailed focus on national conservation concerns. Ultimately, we reviewed the representation of total and priority (i.e. threatened and endemic) taxa in legislation annexes to provide a baseline for unbiased, taxonomy-informed revisions to foster more ambitious actions by Italian stakeholders.

## 2. Methods

### 2.1. Data compilation

Italian checklists of fauna, flora, fungi, lichens and algae were collected from published literature or databases managed by Italian institutions and initiatives. The faunal list of taxa was based on the original Checklist of Italian Fauna, supplemented by the New Checklist of Italian Fauna by LifeWatch Italy (Bologna et al., 2022). Portions published as data papers and applied a class-level cut-off for the inclusion were included (see Supplementary References). The only exception to this threshold was represented by the family Simuliidae (Diptera). Faunal checklists omitted explicit information on vagrancy and irregular breeding status for bird taxa. Therefore, we used the CISO-COI checklist of Italian birds (Baccetti and Fracasso, 2020) to exclude vagrant and irregularly breeding taxa, which are generally assessed at the national level. Alien taxa, when present, were removed. Finally, checklists for Arachnida and Trichoptera were obtained from the Araneae.it and Trichoptera.it web pages. Unlike faunal taxa, checklists of floral and algal species were retrieved from primary literature, whereas the compilation of fungal and lichen taxa relied on contributions from the curators of the Project Dryades (https://dryades.units.it/home/?procedure=checklists). Each checklist was curated to include information about endemism status, although not available for algal and lichen taxa. Where notes about subendemic populations were given, these were resolved as “non-endemic”.

For the IUCN lists at Italian level, these were sourced from the Italian IUCN Committee and published literature. Usually the European, Mediterranean and global red list assessments are freely accessible at the IUCN API at iucnredlist.org by using the advanced search of the website and filtering for the “Global”, “Mediterranean” and “European” regional assessments and “Italy” in the “Land Region” filter (accessed in March 2025). In addition, every European red list published on the European Commission website (https://environment.ec.europa.eu/topics/nature-and-biodiversity/european-red-list-threatened-species_en) was downloaded. Lastly, the annex lists of the Birds and Habitats Directives, the Convention on the Conservation of European Wildlife and Natural Habitats (hereafter Bern Convention), the Convention for the Protection of the Mediterranean Sea Against Pollution (Barcelona Convention) and the Convention on International Trade in Endangered Species of Wild Fauna and Flora (CITES)were obtained by searching the related relative institutional websites of the European Nature Information System (EUNIS, https://eunis.eea.europa.eu/index.jsp) (Detailed references of checklists, red lists and primary literature can be found in Supplementary References).

### 2.2 Taxonomic harmonization and data processing

Scientific names in all documents gathered were harmonized to eliminate cases of synonymy (Pyle, 2016; Grenié et al., 2023). While the intent was not to tamper with the taxonomies in the source material, this was necessary to reduce the variability and obsolesce of older sources and merge each taxon correctly. For this reason, the packages *taxize* (Chamberlain and Szöcs, 2013) in RStudio 4.5.1 (Posit Team, 2025) were employed to resolve synonyms and extract the accepted names for each taxon from global checklists. Similarly, the R package *rgbif (*Chamberlain et al., 2012) was used to extract the most recent names of classes, orders and families. Harmonized datasets where then merged using the package *dplyr* (Wickham et al., 2023) and pivoted tables for visualizations were created with *tidyr* (Wickham et al., 2025b). To ensure consistency with national red list assessments, infra-specific taxa (subspecies, varieties, and forms) were included in the analysis rather than being aggregated at the species level. For many entries in the checklists, harmonization returned the species rank as accepted name. If this occurred for subtaxa whose assessment was available, we retained the original name to maintain taxonomic granularity. Conversely, subtaxa lacking individual assessment were imputed with the status of their respective species. We also resolved homotypic synonyms and addressed nomenclatural homonyms to ensure each taxon was represented by a single unique identifier. We excluded 395 taxa in more taxonomically uncertain groups like insects (i.e. Coleoptera, Lepidoptera, Hymenoptera) and corals. These were assessed in Italian red lists, although we found no correspondence in checklists (details on names and Italian assessments in Supplementary Table 1).

After taxonomic harmonization, IUCN categories for each assessment were then coded assigning a numerical code to each category (from Least Concern to Critically Endangered: 0-5; Possibly Extinct in the Wild: 6; Extinct in the Wild: 7; Possibly Extinct: 8; Regionally Extinct: 9; Extinct: 10) to create an ordinal variable used to compare across the different assessments. Specifically, Data Deficient (DD) taxa were placed at a higher rank than “Least Concern” and “Near Threatened” (hence coded “2”) following a precautionary principle. This is based on recent knowledge that species assigned to Data Deficient classification for lack of data on their distribution might be threatened (Parsons, 2016; Borgelt et al., 2022). The converted threat status was also used to compare threat conditions with European, Mediterranean and global assessments. Given an endemic taxon’s geographical range is restricted within Italian borders (i.e. no other population of assessed taxon is present outside Italy), threat status was assumed to be identical across geographical scopes of each IUCN assessment. However, drawing from the observations of Brito et al. (2010) and Glasnović et al. (2024) on mismatches across red lists, we also investigated possible differences across lists for endemic species by employing the conversion in numbers of each threat status.

Similarly, we coded each taxon per each class/order to indicate inclusion in at least one legislation. All figures were produced using the R package *ggplot2* (Wickham et al., 2025a) and *ggrepel* (Slowikowski, 2016). The results and visualizations follow the non-hierarchical taxonomical format of Italian red list publications to directly compare assessment coverage and biases against full checklists and legislation. Specifically, the invertebrate taxa not included in any list were grouped together (Table 1 for details).

**Table 1:** Total counts with percentages in brackets of major taxa groups assessed in Italian red lists. Percentages for endemics are calculated according to the total of endemic taxa in the “Endemics” column. The group “Other invertebrates” includes all the phyla with no assessment in any list: Acanthocephala, Annelida, Arthropoda, Brachiopoda, Bryozoa, Chaetognatha, Cnidaria (excluding Anthozoa), Ctenophora, Dicyemida, Echinodermata, Entoprocta, Gastrotricha, Gnathostomulida, Hemichordata, Kamptozoa, Kinorhyncha, Loricifera, Nematoda, Nematomorpha, Nemertea, Orthonectida, Phoronida, Platyhelminthes, Porifera, Priapulida, Rotifera, Sipuncula, Tardigrada. For Arthropoda, the remaining classes included in this table were Branchiura, Branchiopoda, Chilopoda, Cirripedia, Diplopoda, Mystacocarida, Ostracoda, Pauropoda, Pentastomida, Pycnogonida, Symphyla and other hexapods in the orders Anthoathecata, Diplura, Entomobryomorpha, Neelipleona, Poduromorpha, Protura and Symphypleona. The nomenclature and taxonomy for each taxon follow https://www.faunaitalia.it/checklist/ *Extinct regionally and/or declared officially extinct worldwide by IUCN. ** Not assessed in Italy but present in checklist

| *Taxa groups in Italian red lists* | *Totals* | *Endemics* | *Assessed* | *Assessed en-*<br>*demics* | *Threatened* | *Threatened en-*<br>*demics* | *Data defi-*<br>*cient* | *Data deficient*<br>*endemics* | *Extinct\** | *Extinct*<br>*endemics\** |
| --- | --- | --- | --- | --- | --- | --- | --- | --- | --- | --- |
| *Algae* | 529 | 0 | 0 (0%) | 0 (0%) | 0 (0%) ** | 0 (0%) | 0 (0%) | 0 (0%) | 0 | - |
| *Anthozoa (Corals)* | 126 | 32 | 100 (79%) | 25 (79%) | 9 (7%) | 1 (3%) | 63 (50%) | 17 (53%) | 0 | 0 |
| *Insects* | 37,896 | 3,730 | 2,224 (6%) | 157 (4%) | 412 (1%) | 63 (2%) | 287 (1%) | 22 (1%) | 9 (0%) | 0 |
| *Lichens* | 3,275 | 0 | 154 (5%) | - | 80 (2%) | - | 13 (0.4%) | - | 29 (1%) | - |
| *Macrofungi* | 4,457 | 6 | 21 (0.5%) | 0 ** | 16 (0.4%) | 0 ** | 2 (0%) | 0 ** | 0 | 0 |
| *Molluscs* | 2,606 | 554 | 0 (0%) | 0 ** | 0 ** | 0 ** | 0 ** | 0 ** | 0 ** | 0 |
| *Non-vascular*<br>*plants* | 1,274 | 0 | 1,257 (99%) | - | 330 (26%) | - | 133 (10%) | - | 23 (2%) | - |
| *Other*<br>*invertebrates* | 16,542 | 2,212 | 0 | 0 ** | 0 ** | 0 ** | 0 ** | 0 ** | - | - |
| *Vascular plants* | 9,002 | 1,757 | 2,524 (28%) | 1,421 (81%) | 596 (7%) | 311 (18%) | 424 (5%) | 260 (15%) | 54 (1%) | 14 (1%) |
| *Vertebrates* | 1,138 | 98 | 1,068 (94%) | 97 (99%) | 185 (16%) | 34 (33%) | 114 (10%) | 5 (5%) | 6 (0.5%) | 0 |
| ***Totals*** | **76,845** | **8,389** | **7,346 (10%)** | **1,700 (20%)** | **1,628 (22%)** | **409 (24%)** | **1,036 (14%)** | **304 (18%)** | **121 (2%)** | **14 (0.2%)** |

### 2.3 Statistical analysis

The association between taxon inclusion in legally binding policies and conservation relevance was evaluated using Fisher’s exact test due to limited representation in policies for some taxonomic groups. We constructed contingency tables comparing the number of endemic taxa versus non-endemic taxa included at least in one legislation, and threatened (i.e. Vulnerable, Endangered or Critically Endangered) versus non-threatened taxa, that were included or excluded from any policy annex. Under the expectation that endemic taxa and taxa facing elevated extinction risk should be preferentially prioritized for legal protection, we tested whether their representation in policy annexes was significantly higher than expected by chance.

## 3. Results

### 3.1 Proportions of taxa assessed by Italian red lists

The final dataset included 76,845 taxa (including subspecies), with 8,389 endemics. Conversely, the count of assessed taxa in Italian red lists was 7,349 (10%), the highest in comparison to other assessment scopes (Figure 1: ,Supplementary Table 1). Proportionately, Italian red lists include the highest number of endemic taxa (n = 1,700, 18%), with effort on vascular plants, with 81% of endemic taxa assessed (n = 1,421) (Figure 1, Supplementary Table 1).

**Figure 1:**
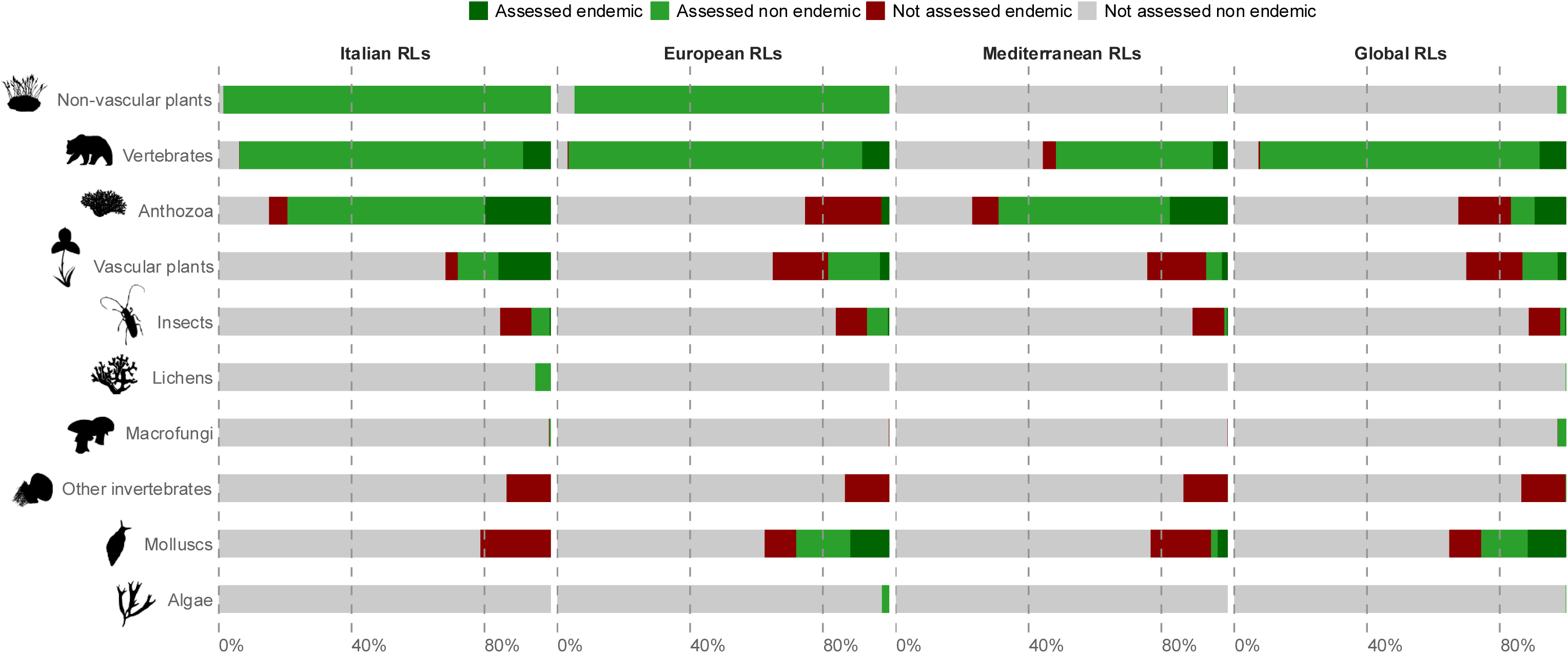
Stacked bar chart displaying the percentage of assessed endemic and non-endemic taxa currently in the Italian red lists, Mediterranean and European lists and global red lists. The silhouettes of each taxon have been resized from phylopic.org.

**Figure 2:**
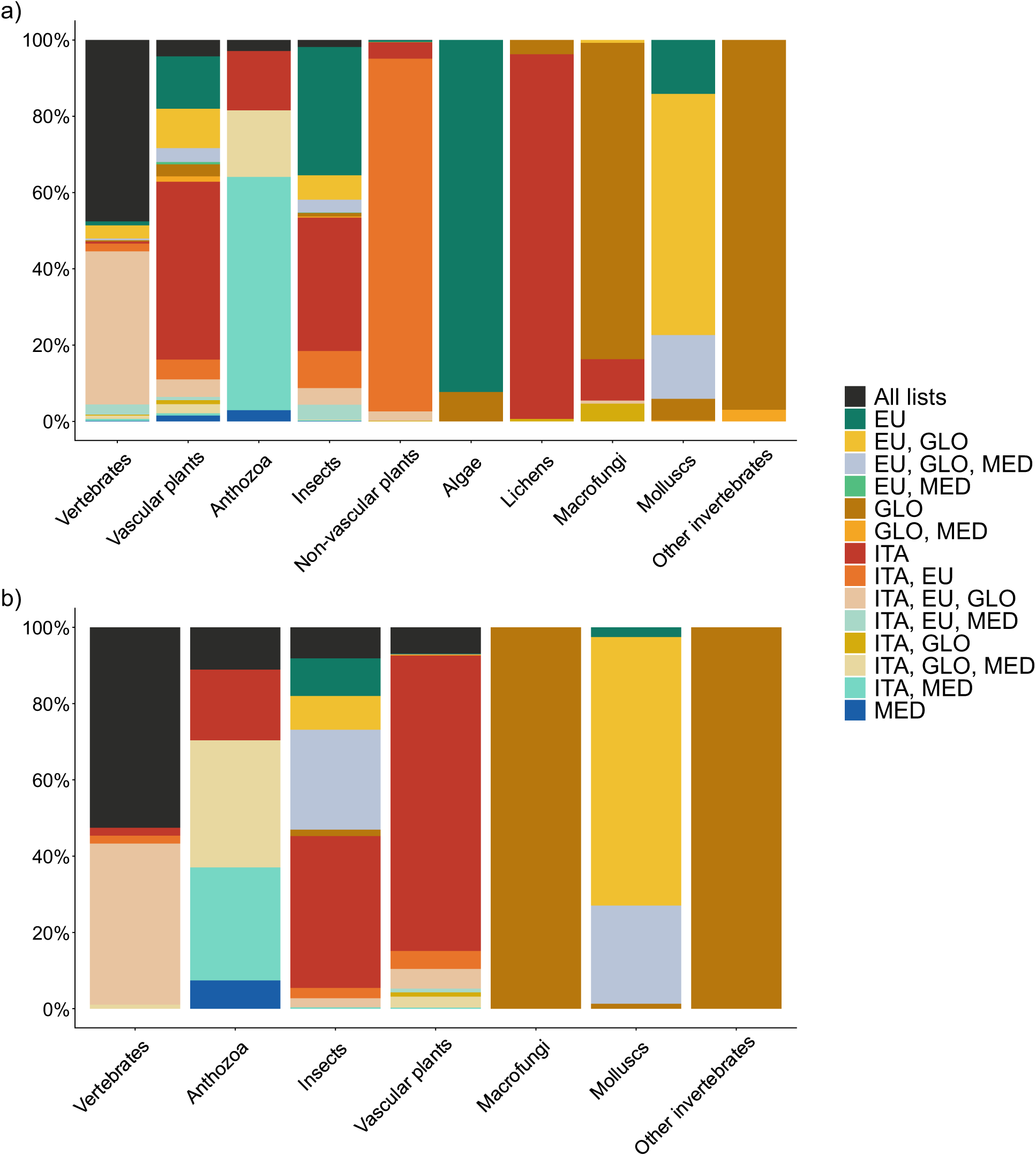
Stacked bar plots showing proportions of assessment coverage in red lists across different geographical scales for a) all taxa and b) endemic-only taxa. The bars have been ordered according to the taxon with highest assessed proportion of taxa in all lists. For graph b), bars show only those macrotaxa in which endemic species are present. Each colour represents presence in a red list geographical scope (present singularly or in more than one): Italian assessment (ITA), European assessment (EU), global assessment (GLO) and Mediterranean assessment (MED).

In Italian assessments, vertebrates are the group with the highest proportion of assessed taxa with 94% of taxa assessed, while all other major taxa are lacking assessments, with the only exception being endemic plants, with more than 80% of them assessed at national level (Figure 1, Table 1). In contrast, insects in Italy remain under-represented, with an assessment rate of only 6%. Odonata are a notable exception with near-complete assessment coverage (87%). They are followed by Coleoptera (13%) and Lepidoptera (5%), while only 2% of Hymenoptera were included, while all remaining orders lacked any national assessment. Corals (Anthozoa) are the only other invertebrate taxa assessed in Italian red lists and to near completion (approximately 80% for total abundance and endemics). Full assessment coverage was achieved for all endemic Odonata and Agnatha, as well as Amphibia, Actinopterygii and Pinopsida. However, it should be noted that these groups possess low richness of endemic taxa. Similarly, high national coverage was recorded for endemic Liliopsida (91%), endemic Magnoliopsida (79%) and endemic corals (78%).

### 3.2 Comparisons across lists

Compared to other assessments, Italian assessments of vascular plants are the most comprehensive (n = 2,524). Nonetheless, non-vascular plants are those for which the number of assessed taxa almost matches the full diversity of the phylum (99%) (Figure 1) but it is scarcely considered at higher geographical scales, with even global lists only including 3% of total richness. For vascular plants, the counts for endemics were up to nine times higher when compared to the Mediterranean list (n = 1,800 in total across all taxa). Lichens have a higher representation in Italian lists (4.7%), twenty-two times higher than the global red list, the only list containing any lichen species (n = 7). Conversely, no Italian assessments were available for algal taxa, although some assessments at European level are found for the phylum Charophyta (12 in Europe, 1 in global lists). Vertebrates are usually almost fully assessed in every list, with Mediterranean red lists including the lowest account (53%). For insects, European assessments are the most complete including 2,553 taxa. This is due to mainly four red lists covering the following orders: Diptera (Syrphidae, n = 414, 6% of totals in order), Hymenoptera (n = 1,028, 13%), Odonata (n = 91, 96%) and Orthoptera (n = 364, 89%). However, Italian red lists better cover the order Coleoptera, with 1,749 taxa assessed. The European red list also provides a coverage of assessment of Italian endemic orthopterans (91% assessed) and for the phylum Mollusca (28% assessed). Exclusively, global red lists contained assessments for all mantids in Italy (n=10) and cephalopods (n = 39), and it is the only list where fragmentary assessments of arachnids (n = 2), copepods (n = 2), holothuroids (n = 6) and malacostracans (n = 23) found in Italy are included. Within the fungal and plant kingdoms, global assessments provide the most extensive coverage for Basidiomycota (n = 114, 3% of total) and Gnetophyta (n = 6, 67%). Finally, while Mediterranean assessments only cover 2% of all taxa in our checklist, they contain assessments for 69% of corals.

### 3.3 Comparison and mismatches of category of status across multi-scale assessments

A comparison of assessment outcomes revealed disparities in extinction risk levels across different geographical scales. Italian red lists assigned a more severe threat category to 35% of taxa compared to European and global lists (Figure 3a, c). In Mediterranean assessments, this percentage lowers to 20%, despite it is to be contextualized in relation to the lower number of taxa assessed.

**Figure 3:**
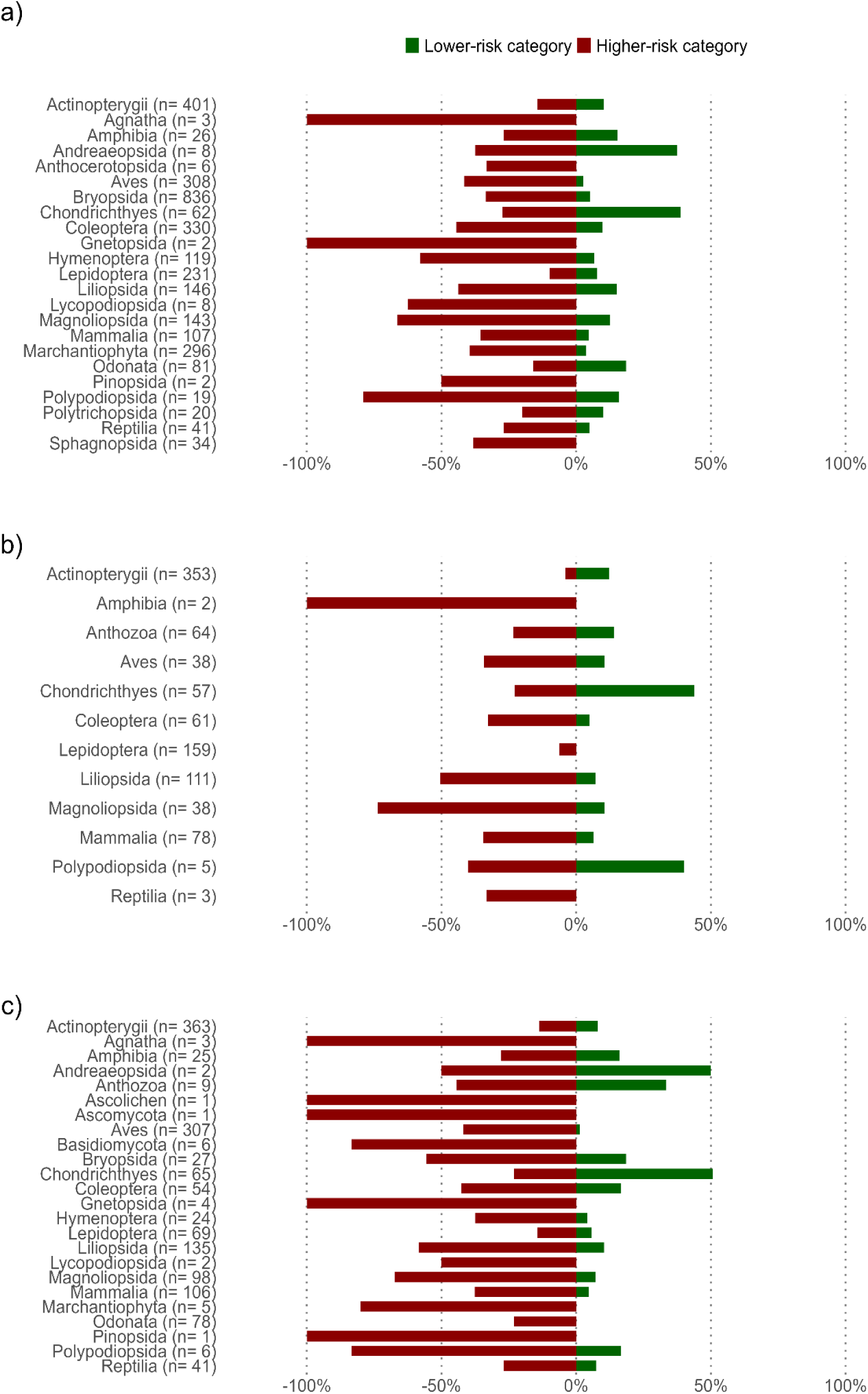
divergent plots showing comparisons of Italian assessments to the other three geographical regions a) Europe, b) Mediterranean and c) globally. The comparisons are expressed as percentage of status over the number of taxa assessed per group. Endemics were excluded as for this graph it is assumed that all endemics have equal assessment. The red bars indicate all taxa whose threat status is more severe (i.e. belong to a higher-risk category in Italian assessments) than the compared assessment, while greens bars are for those with a threat status less severe. Non-endemic taxa with the same assessment were excluded from this representation.

Conversely, a small portion of Italian taxa was found to be in better condition nationally than in Mediterranean (10%) or European or global (8%) lists (Figure 3b). Taxonomic groupings with highest percentages of taxa in higher threat categories are also those with high taxonomic specificity such as Agnatha, Anthocerotopsida, Gnetopsida, Lycopodiopsida and Pinopsida in both European and global lists (Figure 3a, c). Notably, Chondrichthyes was the only class consistently showing a better conservation status at the national level across all three comparative assessments. Italian red lists also include the highest counts of Data Deficient taxa (Figure 4).

**Figure 4:**
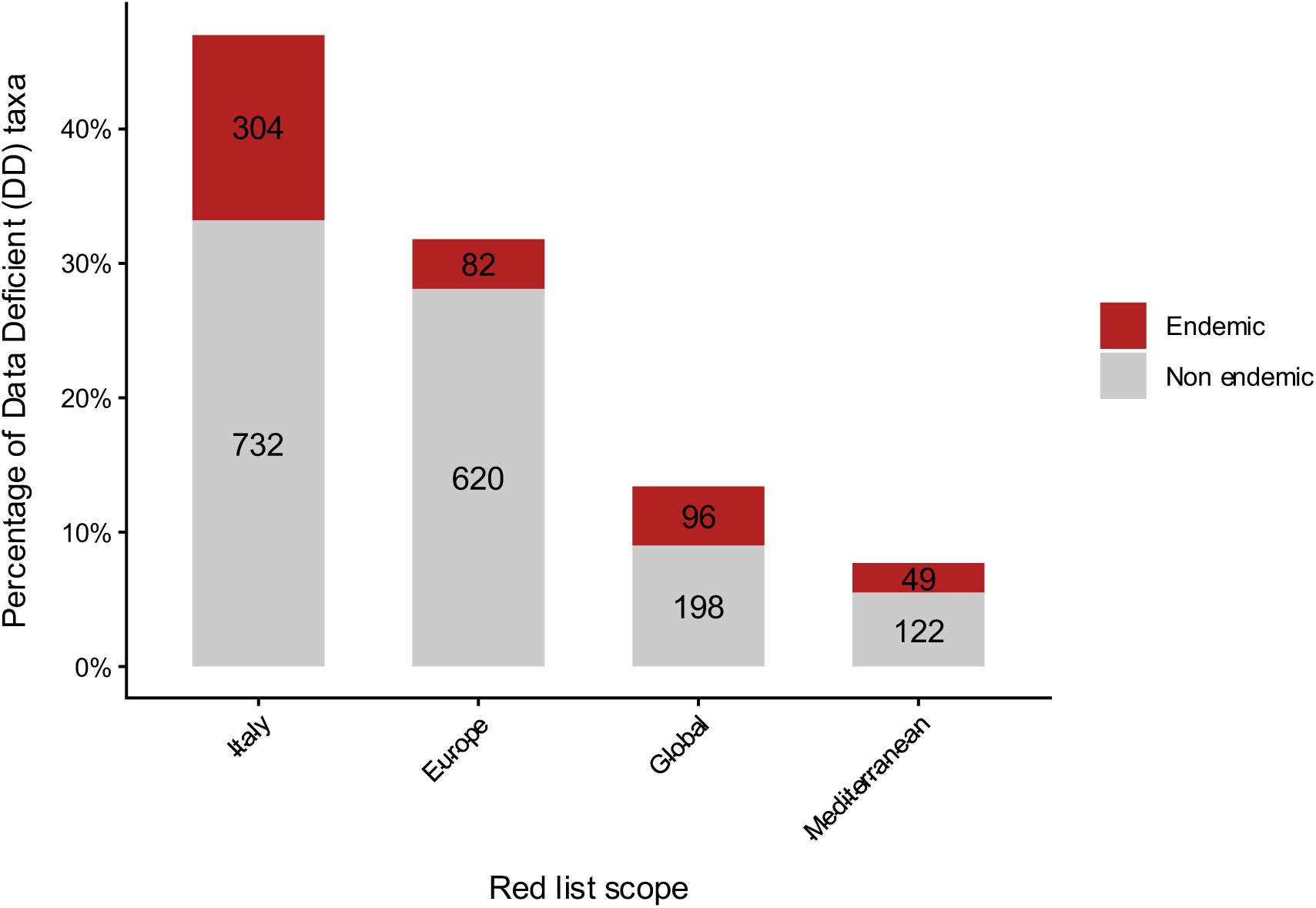
Counts of Data Deficient taxa across IUCN red list geographical scopes.

Mismatching assessments were observed for endemic taxa, even when accounting for the number of taxa whose assessment fell under the species nomenclature (n = 78). Overall, 218 endemics were assigned to different categories across lists, with plant taxa in classes Magnoliopsida and Liliopsida showing the highest count (Figure 5).

**Figure 5:**
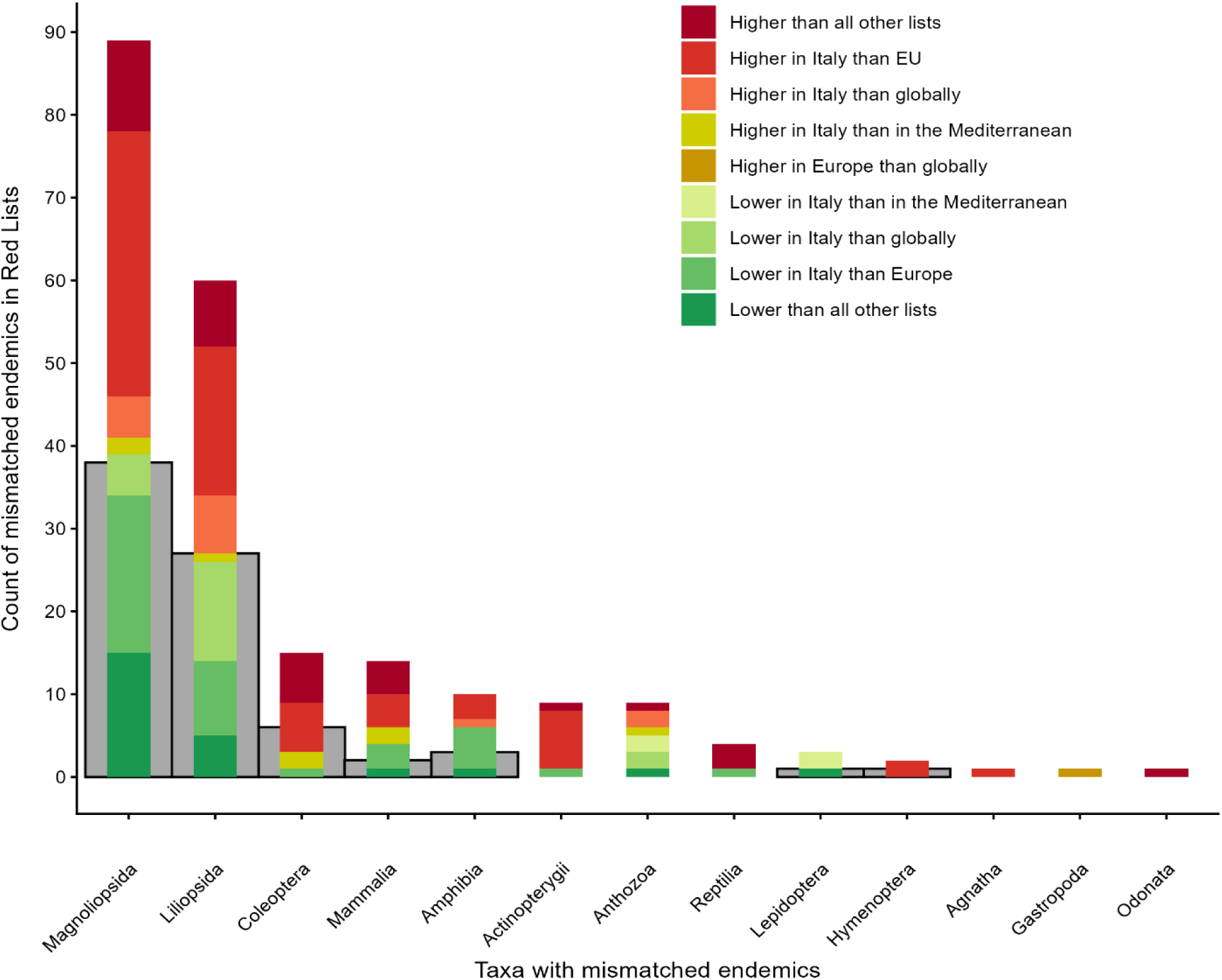
Counts of endemic taxa per each major taxa group with its assessing differing across one or more assessments (coloured bars). The grey bars display the number of subspecies under a species assessment.

### 3.4 Legal protection of Italian taxa and endemics

In total, 1,346 taxa are protected in at least one form of legislation, equal to the 2% of all examined taxa. For endemic taxa, the proportion of protected species is marginally higher at 3% (n = 226). Vertebrates cover half of the total protected taxa and occupy more than 70% of entire of protected endemics. Taxa included in each legislation range between 882 in the Birds and Habitats Directives and 147 in the Barcelona Convention, although the latter has a focus on marine taxa only. Conversely, the Bern Convention and CITES list 812 and 423 taxa respectively. While Birds and Habitats Directives protect nearly half of all vertebrates (47%), average legal coverage for all other groups remains below 4%. The Bern Convention showed a similar patter with 51% vertebrates included. Corals are instead the group with the highest inclusion in the Barcelona appendices (9%) and in CITES (25%).

Fisher’s exact test results revealed that legal inclusion for most taxonomic groups is independent of both threatened and endemic status. While threatened status significantly increased the likelihood of protection for Actinopterygii, Chondrichthyes, Lepidoptera, and Coleoptera, a negative association was observed for ascolichens (Supplementary Table 4a). Endemic status showed significant associations in Actinopterygii, Liliopsida, Magnoliopsida, Anthozoa, and Lepidoptera (Supplementary Table 4b). Endemic fish and plants exhibited increased inclusion likelihood, whereas Anthozoa and Lepidoptera were associated negatively.

As shown in Figure 6a, birds are largely protected, with 99% of threatened and data deficient taxa included in at least one appendix, a trend followed by other vertebrate orders. Amongst insects, Lepidoptera are the order with most threatened taxa in legislations with 7 included over 16 threatened taxa (44%), while Odonata follow with 20% of units included. Coleoptera have the highest number of threatened taxa in the Insecta class (n = 374), however only 2% of these have been included in legislation. For the largest classes of plants – Magnoliopsida and Liliopsida – the percentage of threatened and data deficient taxa in the appendixes is 10% and 27% respectively. The ten Sphagnopsida threatened taxa are all included in a legislation (Habitats Directive), while only two mosses (Bryopsida) out of 231 are protected. Finally, 13% of ascolichens have protection. In contrast, no fungal taxa are protected.

**Figure 6:**
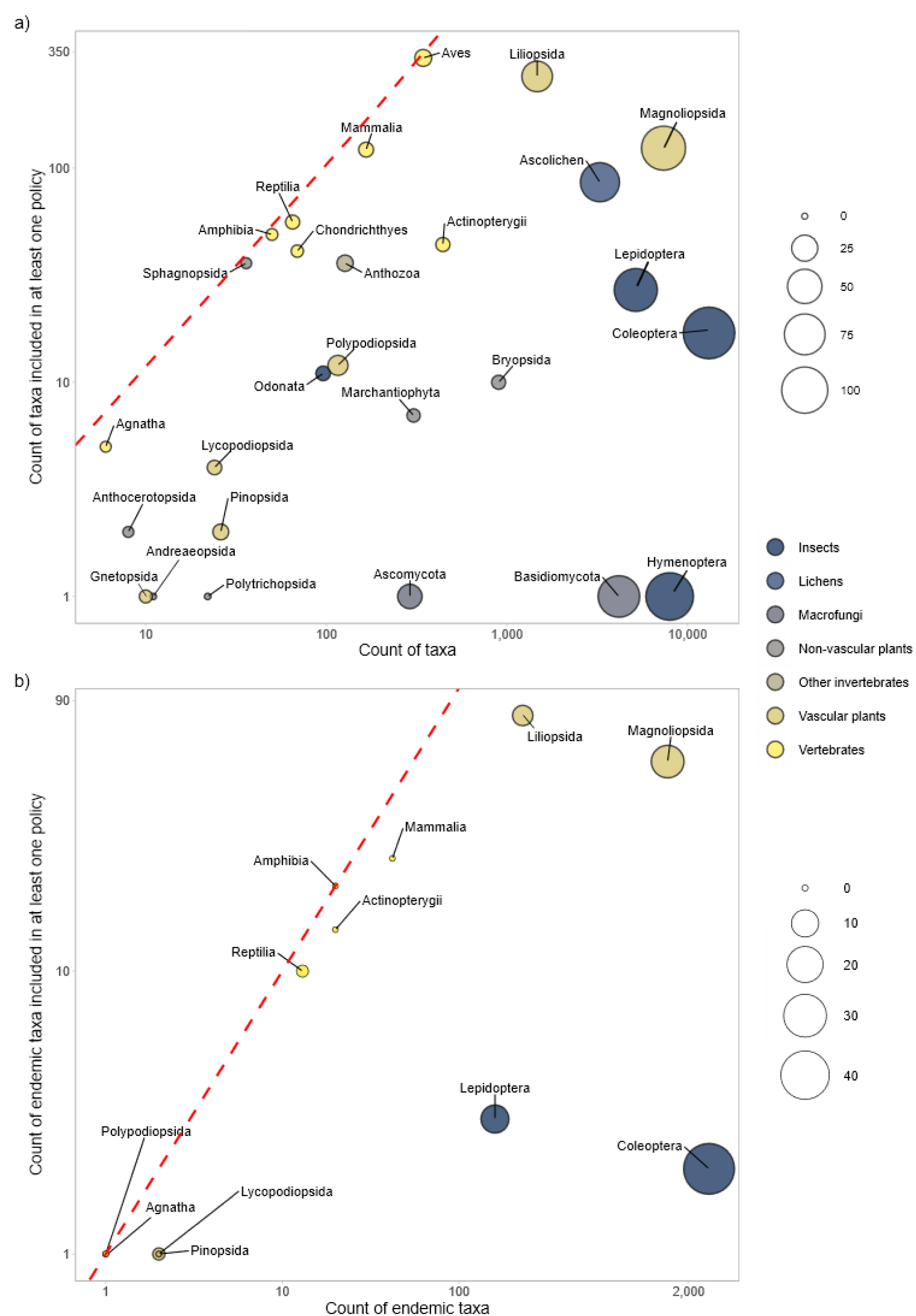
Bubble plots showing the counts for threatened and data deficient **a)** total taxa and **b)** only endemic taxa included in at least one of the policies considered. The colour hues represent the major phyla or divisions, while each bubble indicates lower taxonomical hierarchies. Bubble size portraits the number of not assessed taxa adjusted by its square root to help visualize the richness of each order or class and avoid graphical bias. The dashed red line represents ideal completeness (100% of threatened taxa covered). Points along or close to the line indicate a comprehensive policy coverage, while those far apart show gaps in coverage.

The protection of endemic biodiversity remains highly fragmented across taxonomic groups (Figure 6b). Lepidoptera exhibit a drop in representation, with only 3 protected endemics over 160 taxa (2%). Mammals also demonstrate a notable inclusion gap, with only 60% of endemic units under protection. Only two endemic Coleoptera taxa are in the appendices (0.1%), and for Magnoliopsida only 4% is protected despite the group’s high richness. Hyper-diverse groups such as Diptera, Orthoptera and other insect orders register no inclusion of Italian endemics under protection. A similar pattern of neglect is observed for Anthozoans, with only 3 out of 32 taxa protected, and for non-insect invertebrates, where a mere 6 out of 2,212 endemic taxa are legally recognized. Conversely, a handful of groups with low absolute endemic richness - specifically Agnatha, Amphibia, Odonata, and ferns (Polypodiopsida) achieve full inclusion in at least one policy framework.

### 3.5 Exclusion of taxa

Using Italian assessments as reference, there were in total 1,321 threatened (Vulnerable, Endangered, Critically Endangered), 55 Possibly Extinct and 36 Extinct in Italy taxa excluded from policies, with net lack of inclusion for Magnoliopsida (n = 447), Coleoptera (n = 368) and Bryopsida (n = 244) compared to other groups (see Figure 7 for categories with the most taxa excluded).Actinopterygii were the only vertebrate group with a substantial number of taxa in threatened categories not listed in policies (n = 15).

**Figure 7:**
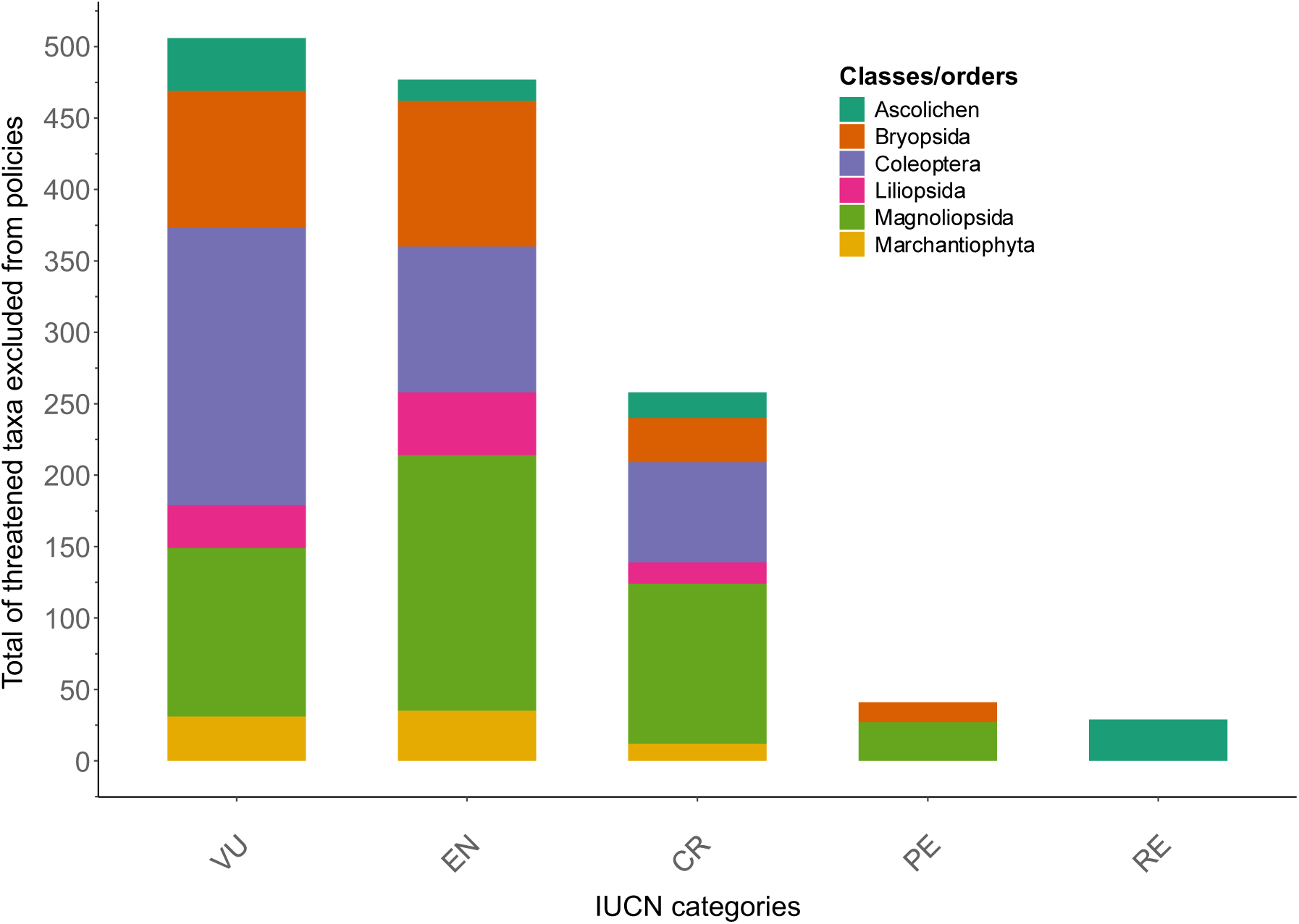
IUCN categories with more than 10 species/subspecies excluded from policies.

## 4. Discussion

### 4.1 Taxonomic bias and impediment in Italian red lists

This first, comprehensive account of Italian biodiversity, red lists and European policies exposes taxonomic biases and impediments, an established dilemma in prioritization and conservation of species (McKinney, 1999; Cardoso et al., 2011a, 2011b; Farooq et al., 2026). Only 10% of Italian taxa across plants, animals, lichens, fungi and algae have been assessed in Italian lists, and only 14% of these are protected at least by one international policy. While Italian red lists are the most extensive lists compared to lists with wider geographical scope, the taxonomic coverage is fragmented, and European and global assessments compensate for the gaps for some phyla (e.g. Mollusca), orders (e.g. Orthoptera) and even kingdoms (Fungi).

As expected, taxonomic bias is evident across Italian lists (Figure 1, Table 1), with invertebrate underrepresentation at concerning levels comparable to recent results in the Neotropics (Justino et al., 2026). Vertebrates are overrepresented in scientific research and funding due to a longer tradition in data availability, charisma and attractiveness (Bonnet et al., 2002; Brambilla et al., 2013; Titley et al., 2017; Mammola et al., 2020; Wang et al., 2021; Fischer et al., 2025; Guénard et al., 2025). Exceptionally, corals are included in high proportions probably favoured by research urgency and favourable public opinion due to climate-driven extinction risk (Figure 2) (Carpenter et al., 2008).

On the contrary, comprehensive assessments for hyper-diverse groups of invertebrates, like Mollusca, still lack, despite them being equally important for ecosystems and characterized by higher rates of extinction than vertebrates (Cowie et al., 2022; Johnson et al., 2025). Comparable taxonomic neglect extends beyond invertebrates. Verlaque et al. (2019) criticize the shortcomings of both assessments and environmental policies for macroalgal diversity by relating them to systematic preferences in terrestrial taxa. This fallacy is attributed to sampling remoteness or, as in the case of fungal species, to complex life cycles that are in contrast with the requirements of conventional IUCN guidelines for listing (Dahlberg and Mueller, 2011). Conservation of Italian fungi relies on assessments published at global level despite an attempt was made to assess a few species to complement the assessment of policy plant species. This suggests that taxonomic updates and coordinated initiatives across the country for data collection in the field are needed to publish a larger assessment for Italian macrofungi. Bryophytes are another phylum subjected to deficit in research and funding (Deilmann et al., 2024). Nonetheless, Italian and European assessments almost fully match the extent of this taxon diversity in Italy. Indeed, the Italian checklist of bryophytes is one of the most recent (Aleffi et al., 2023), showing that the Italian bryological community opts for publication of updated taxonomy and assessments rigorously.

Full assessment coverage of other hyper-diverse groups like insects is one of the greatest challenges for IUCN due to their diversity and rate of description of new species. Insects are one of the most assessed classes in Italy despite taxonomic impediments. For instance, Italian lists rank first in number of Coleoptera assessed (see Supplementary Table 2) in the lists here considered, associating an adequate conservation attention to this order, despite expertise in coleopteran identification and description has been assigned to the Poor Capacity (PO) category in European Red List of Insect Taxonomists (Hopkins and Freckleton, 2002; Hochkirch et al., 2022). Italian assessments mainly include saproxylic beetles, suggesting that their functional importance in regulating forest ecosystems has pushed assessors to allocate resources for the creation of the list. Resource allocation in assessments and protection depends on feasibility (Wilson et al., 2011; Wiedenfeld et al., 2021). Hence, for large taxonomic groups, assessors may prioritize taxa of functional importance, phylogenetic uniqueness or with aesthetic value instead of aiming to assess their entire diversity comprehensively. Consistently, in Azam et al. (2016), Italy’s national red lists demonstrated preferences towards well-known taxa. Furthermore, assessors are motivated to evaluate only those species with distribution data or resort to personal knowledge of historical versus current distribution to assign extinction status (Tomasini, 2018). In Italy, well-known taxa such as all vertebrate classes and insects like Odonata have kept the pace with counts observable in Europe (Azam et al., 2016). By contrast, Lepidoptera, despite being a popular order amongst assessors, due to better data availability and taxonomic knowledge, was not covered as largely in Italy, likely due to a wider proportion of less-charismatic taxa (e.g. micromoths). Lastly, the limited representation of Hymenoptera in Italian red lists exemplifies how motivation to assess such functionally important taxa is constrained by a lack of fundamental biological, taxonomic and distributional data (Marshall et al., 2024).

### 4.2 The fine-scale scope of national red lists: endemics, Data Deficient taxa and subspecies

Our analysis reveals the importance of Italian red lists in publishing novel assessments for endemics, effectively exposing conservation gaps previously only identified in Sicily (Di Gristina, et al., 2022). Such multiscale comparisons help highlight the importance of national red lists to catalyze the publication of endemic species assessments in the main global list when no regional assessments exists (Glasnović et al., 2024), in turn leading to larger funding and public interest (Azam et al., 2016; Karam-Gemael et al., 2018, 2020; Barahona-Segovia and Zúñiga-Reinoso, 2021). Interestingly, when endemics were assessed across scales, several of these assessments did not match (Figure 5). Beyond cases attributable to the referencing of subspecific taxa under their parent taxon (Figure 5), discrepancies in assessments of endemic taxa across geographical levels genuinely reflect systematic inconsistencies in following assessment protocols. We hypothesize that these mismatches are caused by co-occurring factors such as lack of coordination amongst teams of assessors in regional and global lists, as well as different taxonomic treatments and distributional data employed (Hilton-Taylor et al., 2000; Brito et al., 2010; Moon et al., 2023; Glasnović et al., 2024). Assuming national assessors employ much granular observational data for their calculations, national lists have the added potential to flag discrepancies, increasing reliability of global lists especially when correcting listing at severe extinction risk categories.

National assessments also provide the high-resolution data necessary to identify localized declines in otherwise widespread taxa (Figure 3). Similarly, we demonstrate the concern of data deficiency at national level, which could trigger a call-to-action to intensify data collection for neglected taxa. This level of granularity is crucial for detecting the destabilization of sink-source dynamics, allowing for more effective threat triage and the prioritization of viable populations that broader, coarse-scale assessments frequently overlook (Azam et al., 2016).

The most striking difference found in Italian lists was high taxonomic resolution given the inclusion of numerous subspecies and newly discovered species while being absent in other assessments. Taxonomic inflation is another considerable impediment to IUCN red list assessments (i.e. the description of new species beyond necessary; Raposo et al., 2021), and we observed that at larger scales assessors may publish assessments that guarantee conservation feasibility over taxonomic refinement. While the inclusion of subspecies carries the acknowledged risk of diverting conservation resources toward inbred and isolated subpopulations when molecular validation is doubtful (Clavero et al., 2024), the benefits are demonstrable where subspecific elevation to species rank reflects genuine evolutionary distinctiveness. For example, the *Dryomys aspromontis* (Bisconti et al., 2018), previously a Calabrian subspecies of *Dryomys nitedula* and recently been upgraded in Italian mammal checklists as species, is only assessed in Italy as recognized amongst Italian taxonomists. Italian red lists therefore prove their potential to reduce taxonomic inaccuracy in assessments, which may result in promotion of data collection schemes for newly described species.

### 4.3 Disparities between Italian red lists and inclusion in policies

The inclusion in the annexes of European Directives, Bern and Barcelona treaties and CITES appendices was expectedly biased towards more charismatic taxa. A promising exception is characterized by Sphagnopsida, a class of mosses comprising keystone species for peatland ecosystems. Similarly to previous literature analysing policy gaps (Cardoso, 2012; Leandro et al., 2017; Mammides, 2019), not only insects are widely overlooked but Italy acts as case study for lack of meaningful inclusion across non-vertebrate diversity. In contrast to the UK, where threatened Lepidoptera are overrepresented in local policies (Duffus and Morimoto, 2022), functionally important insect taxa like Hymenoptera, Coleoptera and Lepidoptera in Italy are severely underrepresented. Notably, while Lepidoptera receive the highest legislative coverage among Italian insects, this inclusion remains insufficient, particularly for the country’s significant endemic diversity. As the European Union is organized differently from the UK with directives required to include species from each communitarian country, it is likely that Italian lepidopteran biodiversity is subjected to a geographical bias in multi-state policies.

By quantifying the fraction of species assessed in national red lists and the even lower percentage then included in policies, we argue that the technical and legal frameworks currently governing decisions for Italian biodiversity create a double bottleneck in decision making, where only a fraction of taxa reach policymakers’ attention. This disparity is consequential if contextualized within the objectives of the Italian National Biodiversity Strategy to 2030. While the Strategy aims to restore healthy populations for 30% of the species in both Directives and 50% of species in national red lists threatened by invasive species, this ambition is significantly skewed due to inclusion biases. By tethering restoration targets primarily to species already listed in Directives, the Strategy risks to reiterate taxonomic biases in funding allocation, in turn reducing the effectiveness of its ambitions. Given their current overrepresentation in these legal frameworks, resources may be disproportionately funnelled toward already well-funded groups like birds and mammals, effectively overlooking the majority of Italy’s threatened invertebrate groups and the near totality of algal and fungal diversity

In this scenario, we suggest two improvements in Italy’s strategy. First, we argue for the adoption of an unbiased score-based prioritization for threatened and endemic taxa, drawing upon the concept of national responsibility, and the publication of a national priority list that complements the inertial adoption of priorities set by multi-state policies (Schmeller et al., 2008a, 2008c). Second, we showed the potential of national red lists to be a valid support to global lists, and we support the fast-tracking the production of national red lists, with data from standardized and consistent data collection campaigns. New methodologies like predictive models with machine learning are also an option to extrapolate extinction risk in data deficient taxa (Borgelt et al., 2022). Beyond endemic status and restricted geographical range, criteria for prioritization should advantage habitat-building taxa such as macroalgae and indicator species such as caddisflies (Trichoptera), so far overlooked even at global levels (Johnson et al., 2025).

We recognize that full coverage of assessments is yet not feasible due to shortcomings in IUCN protocols when applied to non-vertebrate groups (e.g. Cardoso et al., 2011a), despite Italy’s strategy includes actions that safeguard such groups still hard to assess. We acknowledge that Italy is aiming to guarantee ecosystem-based protection and connectivity among protected areas to ensure long-term protection of interspecific processes, genetic diversity and buffer climate range shifts even for taxa with lower extinction concern (Pryke and Samways, 2012; Attorre et al., 2018; Moat et al., 2019). In addition, 172 Key Biodiversity Areas (KBA) are being developed as sites with high importance for total biodiversity and for specific taxa (Kullberg et al., 2019; Nania et al., 2024a, 2024b).

Collectively, we recognize that the patterns documented are not unique to Italy. Vertebrate overrepresentation and invertebrate neglect are recurring hardships of conservation protection legislation and reporting worldwide, and the taxonomic biases we identify reflect tendencies embedded in how assessors and policymakers allocate attention and resources. The Italian case however differs in the convergence of high endemism, a central position within the Mediterranean biodiversity hotspot, and a renewed momentum for environmental conservation provided by the Nature Restoration Regulation. Together, these factors make Italy a useful case to criticize how red lists and environmental policies have been so far compiled. Realizing this potential requires conservation strategies that minimize the taxonomic and observational biases that, left unaddressed, will continue to distort prioritization regardless of the ambition of the frameworks they underpin.

## Supporting information

Supplementary References

