## Supplementary References for "How well is Italian biodiversity represented in red lists and conservation legislation? Taxonomic biases, coverage gaps and the assessment-to-legislation bottleneck"

### Checklists


1) Aleffi, M., Cogoni, A. & Poponessi, S., 2023. An updated checklist of the bryophytes of Italy, including the Republic of San Marino and Vatican City State. *Plant Biosystems – An International Journal Dealing with all Aspects of Plant Biology*, 157(6), pp.1259–1307. https://doi.org/10.1080/11263504.2023.2284136
2) Angelini, P. et al., 2017. Current knowledge of Umbrian macrofungi (central Italy). *Plant Biosystems – An International Journal Dealing with all Aspects of Plant Biology*, 151(5), pp.915–923. https://doi.org/10.1080/11263504.2016.1265609
3) Baccetti, N. & Fracasso, G., 2021. CISO-COI Checklist of Italian birds-2020. *Avocetta*, 45(1). https://doi.org/10.30456/avo.2021_checklist_en
4) Bartolucci, F. et al., 2024. A second update to the checklist of the vascular flora native to Italy. *Plant Biosystems – An International Journal Dealing with all Aspects of Plant Biology*, 158(2), pp.219–296. https://doi.org/10.1080/11263504.2024.2320126
5) Becker, R., 2019. The Characeae (Charales, Charophyceae) of Sardinia (Italy): habitats, distribution and conservation. *Webbia*, 74(1), pp.83–101. https://doi.org/10.1080/00837792.2019.1607497
6) Bologna, M.A., Boero, F., Bonato, L., Casale, A., Curini Galletti, M., Massa, B., Minelli, A., Oliverio, M., Vigna Taglianti, A. & Zapparoli, M., 2018. *The new Checklist of the Italian fauna*. In: *Conferenza Annuale LifeWatch Italia 2018*, 5–27 June 2018.
7) Ciadamidaro, S. & Mancini, L., 2022. The new Checklist of the Italian Fauna: Simuliidae. *Biogeographia – The Journal of Integrative Biogeography*, 37(2). http://dx.doi.org/10.21426/B637257479
8) Cianferoni, F., Carapezza, A. & Faraci, F., 2024. The new Checklist of the Italian Fauna: Heteroptera. *Biogeographia – The Journal of Integrative Biogeography*, 39(2). https://doi.org/10.21426/B639263688
9) Ferraro, V. et al., 2022. The checklist of Sicilian macrofungi. *Journal of Fungi*, 8(6), 566. https://doi.org/10.3390/jof8060566
10) La Porta, G. et al., 2023. The new checklist of the Italian fauna: Odonata. *Biogeographia – The Journal of Integrative Biogeography*, 38(1). https://doi.org/10.21426/B638158781
11) Letardi, A., 2022. The new checklist of the Italian fauna: Mecoptera. *Biogeographia – The Journal of Integrative Biogeography*, 37(1). https://doi.org/10.21426/B637156506
12) Lodovici, O. & Valle, M., 2020. *Checklist dei Tricotteri Italiani*. Bergamo: Museo Civico Scienze Naturali Enrico Caffi.
13) Minelli, A. & Stoch, F., 2007. The checklist of the Italian fauna. In: Ruffo, S. & Stoch, F. (eds.), *Checklist and distribution of the Italian fauna*, vol. 10, pp.21–23.
14) Nimis, P.L., 2025. ITALIC – The Information System on Italian Lichens. Version 8.0. University of Trieste, Dept. of Biology. Available at: https://dryades.units.it/home/index.php (Accessed 28 June 2025).
15) Pagana, I. et al., 2024. New Characeae (Charophyceae, Charales) report in eastern Sicily (Italy). *Italian Botanist*, 18, pp.109–122. https://doi.org/10.3897/italianbotanist.18.131661
16) Pantini, P. & Isaia, M., 2019. Araneae.it: the online catalog of Italian spiders, with addenda on other arachnid orders occurring in Italy (Arachnida: Araneae, Opiliones, Palpigradi, Pseudoscorpionida, Scorpiones, Solifugae). *Fragmenta Entomologica*, 51(2), pp.127–152. https://doi.org/10.13133/2284-4880/374

17) Relini, G. (ed.), 2008, *Checklist della flora e della fauna dei mari italiani / Checklist of the flora and fauna in Italian seas. Parte II.*
18) Renda, W. et al., 2022. The new checklist of the Italian fauna: marine Mollusca. *Biogeographia – The Journal of Integrative Biogeography*, 37(1). https://doi.org/10.21426/B637156028
19) Schifani, E., 2022. The new checklist of the Italian fauna: Formicidae. *Biogeographia – The Journal of Integrative Biogeography*, 37(1). https://doi.org/10.21426/B637155803

### Italian Red Lists

20) Audisio, P., Baviera, C., Carpaneto, G.M., Biscaccianti, A.B., Battistoni, A., Teofili, C. & Rondinini, C. (compilatori), 2014. *Lista Rossa IUCN dei coleotteri saproxilici italiani*. Comitato Italiano IUCN and Ministero dell'Ambiente e della Tutela del Territorio e del Mare, Rome.
21) Balletto, E., Bonelli, S., Barbero, F., Casacci, L.P., Sbordoni, V., Dapporto, L., Scalercio, S., Zilli, A., Battistoni, A., Teofili, C. & Rondinini, C. (compilatori), 2015. *Lista Rossa IUCN delle farfalle italiane – Ropaloceri*. Comitato Italiano IUCN and Ministero dell'Ambiente e della Tutela del Territorio e del Mare, Rome.
22) Gheza, G., Nascimbene, J., Lelli, C., Benesperi, R., Giordani, P., Paoli, L. & Brunialti, G., 2022. Towards a Red List of the terricolous lichens of Italy. *Plant Biosystems – An International Journal Dealing with all Aspects of Plant Biology*, 156(3), 824–825.
23) Gustin, M., Nardelli, R., Brichetti, P., Battistoni, A., Rondinini, C. & Teofili, C. (compilatori), 2021. *Lista Rossa IUCN degli uccelli nidificanti in Italia 2021*. Comitato Italiano IUCN and Ministero dell'Ambiente e della Tutela del Territorio e del Mare, Rome.
24) Quaranta, M., Cornalba, M., Biella, P., Comba, M., Battistoni, A., Rondinini, C. & Teofili, C. (compilatori), 2018. *Lista Rossa IUCN delle api italiane minacciate*. Comitato Italiano IUCN and Ministero dell'Ambiente e della Tutela del Territorio e del Mare, Rome.
25) Relini, G., Tunesi, L., Vacchi, M., Andaloro, F., D'Onghia, G., Fiorentino, F., Garibaldi, F., Orsi Relini, L., Serena, F., Silvestri, R., Battistoni, A., Teofili, C. & Rondinini, C. (compilatori), 2017. *Lista Rossa IUCN dei pesci ossei marini italiani*. Comitato Italiano IUCN and Ministero dell'Ambiente e della Tutela del Territorio e del Mare, Rome.
26) Rondinini, C., Battistoni, A. & Teofili, C. (compilatori), 2022. *Lista Rossa IUCN dei vertebrati italiani 2022*. Comitato Italiano IUCN and Ministero dell'Ambiente e della Sicurezza Energetica, Rome.
27) Rossi, G., Montagnani, C., Gargano, D., Peruzzi, L., Abeli, T., Ravera, S., Cogoni, A., Fenu, G., Magrini, S., Gennai, M., Foggi, B., Wagensommer, R.P., Venturella, G., Blasi, C., Raimondo, F.M. & Orsenigo, S. (eds.), 2013. *Lista Rossa della Flora Italiana. 1. Policy species e altre specie minacciate*. Comitato Italiano IUCN and Ministero dell'Ambiente e della Tutela del Territorio e del Mare, Rome.
28) Rossi, G., Orsenigo, S., Gargano, D., Montagnani, C., Peruzzi, L., Fenu, G., Abeli, T., Alessandrini, A., Astuti, G., Bacchetta, G., Bartolucci, F., Bernardo, L., Bovio, M., Brullo, S., Carta, A., Castello, M., Cogoni, D., Conti, F., Domina, G., Foggi, B., Gennai, M., Gigante, D., Iberite, M., Lasen, C., Magrini, S., Nicolella, G., Pinna, M.S., Poggio, L., Prosser, F., Santangelo, A., Selvaggi, A., Stinca, A., Tartaglini, N., Troia, A., Villani, M.C., Wagensommer, R.P., Wilhalm, T. & Blasi, C., 2020. *Lista Rossa della Flora Italiana. 2. Endemiti e altre specie minacciate*. Ministero dell'Ambiente e della Tutela del Territorio e del Mare, Rome.
29) Riservato, E., Fabbri, R., Festi, A., Grieco, C., Hardersen, S., Landi, F., Utzeri, C., Rondinini, C., Battistoni, A. & Teofili, C. (compilatori), 2014. *Lista Rossa IUCN delle libellule italiane*. Comitato Italiano IUCN and Ministero dell'Ambiente e della Tutela del Territorio e del Mare, Rome.

30) Salvati, E., Bo, M., Rondinini, C., Battistoni, A. & Teofili, C. (compilatori), 2014. *Lista Rossa IUCN dei coralli italiani*. Comitato Italiano IUCN and Ministero dell'Ambiente e della Tutela del Territorio e del Mare, Rome.

### Global, European and Mediterranean Red Lists

1) Allen, D.J., Bilz, M., Leaman, D.J., Miller, R.M., Timoshyna, A. & Window, J., 2014. *European Red List of Medicinal Plants*. Publications Office of the European Union, Luxembourg. https://doi.org/10.2779/907382
2) Bilz, M., Kell, S.P., Maxted, N. & Lansdown, R.V., 2011. *European Red List of Vascular Plants*. Publications Office of the European Union, Luxembourg. https://doi.org/10.2779/8515
3) BirdLife International, 2021. *European Red List of Birds*. Publications Office of the European Union, Luxembourg.
4) Cálix, M. et al., 2018. *Supplementary material to the IUCN European Red List of Saproxylic Beetles*. IUCN, Brussels.
5) Cox, N.A. & Temple, H.J., 2009. *European Red List of Reptiles*. Office for Official Publications of the European Communities, Luxembourg.
6) Cuttelod, A., Seddon, M. & Neubert, E., 2011. *European Red List of Non-marine Molluscs*. Publications Office of the European Union, Luxembourg.
7) Freyhof, J. & Brooks, E., 2011. *European Red List of Freshwater Fishes*. Publications Office of the European Union, Luxembourg.
8) García Criado, M., Väre, H., Nieto, A., Bento Elias, R., Dyer, R., Ivanenko, Y., Ivanova, D., Lansdown, R., Molina, J.A., Rouhan, G., Rumsey, F., Troia, A., Vrba, J. & Christenhusz, M.J.M., 2017. *European Red List of Lycopods and Ferns*. IUCN, Brussels.
9) Hochkirch, A. et al., 2016. *European Red List of Grasshoppers, Crickets and Bush-crickets*. Publications Office of the European Union, Luxembourg.
10) Hodgetts, N. et al., 2019. *A miniature world in decline: European Red List of Mosses, Liverworts and Hornworts*. IUCN, Brussels. https://doi.org/10.2305/IUCN.CH.2019.ERL.2.en
11) IUCN, 2025. *The IUCN Red List of Threatened Species*. Version 2025–2. Available at: https://www.iucnredlist.org (accessed 2025).
12) Kalkman, V.J. et al., 2010. *European Red List of Dragonflies*. Publications Office of the European Union, Luxembourg.
13) Neubert, E. et al., 2019. *European Red List of Terrestrial Molluscs*. IUCN, Cambridge, UK and Brussels, Belgium. Available at: https://portals.iucn.org/library/node/48439
14) Nieto, A. et al., 2014. *European Red List of Bees*. Publications Office of the European Union, Luxembourg.
15) Numa, C., Tonelli, M., Lobo, J.M., Verdú, J.R., Lumaret, J.-P., Sánchez-Piñero, F., Ruiz, J.L., Dellacasa, M., Ziani, S., Arriaga, A., Cabrero, F., Labidi, I., Barrios, V., Şenyüz, Y. & Anlaş, S., 2020. *The conservation status and distribution of Mediterranean dung beetles*. IUCN, Gland and Málaga.
16) Otero, M.M. et al., 2017. *Overview of the conservation status of Mediterranean anthozoans*. IUCN, Málaga.
17) Rivers, M.C. et al., 2019. *European Red List of Trees*. IUCN, Cambridge, UK and Brussels, Belgium.
18) Schubert, H. et al., 2025. *Charophytes of Europe*. [Monograph]. https://doi.org/10.1007/978-3-031-31898-6

19) Temple, H.J. & Cox, N.A., 2009. *European Red List of Amphibians*. Office for Official Publications of the European Communities, Luxembourg.
20) Temple, H.J. & Terry, A. (compilers), 2007. *The status and distribution of European mammals*. Office for Official Publications of the European Communities, Luxembourg.
21) Van Swaay, C. et al., 2010. *European Red List of Butterflies*. Publications Office of the European Union, Luxembourg.
22) Vujić, A. et al., 2022. *Pollinators on the edge: European Red List of Hoverflies*. European Commission, Brussels.
23) Wilson, B., Beech, E., Window, J., Allen, D.J. & Rivers, M., 2019. *European Red List of selected endemic shrubs*. IUCN, Brussels and Cambridge. Available at: https://portals.iucn.org/library/node/48438

### Legislation annexes

1) Convention on International Trade in Endangered Species of Wild Fauna and Flora (CITES) (1973). Available at: https://checklist.cites.org/#/en. Accessed October 2025.
2) Council of Europe (1979) *Convention on the Conservation of European Wildlife and Natural Habitats*. Document 104, Strasbourg, France.
3) Council of the European Communities (1992) Council Directive 92/43/EEC of 21. May 1992 on the conservation of natural habitats and of wild fauna and flora. Official Journal of the European Communities, **35**, 7–50.
4) European Nature Information System (EUNIS), Available at: https://eunis.eea.europa.eu/. Accessed April 2025.
5) European Commission (1992). Council Directive 79/ 409/EEC of 2 April 1979, on the conservation of wild birds. European Community environment legislation 4: 2-49.
6) UNEP, Barcelona Convention and Protocols. Available at: https://www.unep.org/unepmap/who-we-are/barcelona-convention-and-protocols.
