## Supplementary Table 1 for "How well is Italian biodiversity represented in red lists and conservation legislation? Taxonomic biases, coverage gaps and the assessment-to-legislation bottleneck"

| **Supplementary Table 1:** Account of 395 invertebrate taxa belonging to the Anthozoa, Coleoptera, Lepidoptera and Hymenoptera Italian Red Lists that were not included in the main count proposed in the article due to mismatches in the taxa checklist assembled for the review. | | | | | |
|---|---|---|---|---|---|
| **Order** | **Family** | **Taxon name** | **Category** | **Canonical name** | **Accepted name** |
| Actiniaria | | | | | |
| | Hormathiidae | *Amphianthus dohrnii* | DD | *Amphianthus dohrnii* | *Amphianthus dohrnii* |
| | Sagartiidae | *Sagartia troglodytes* | DD | *Cylista troglodytes* | *Sagartia troglodytes* |
| Alcyonacea | | | | | |
| | Elliselliidae | *Elisella paraplexauroides* | DD | *Elisella paraplexauroides* | *Elisella paraplexauroides* |
| | | *Viminella flagellum* | LC | *Viminella flagellum* | *Viminella flagellum* |
| | Plexauridae | *Spinimuricea klavereni* | DD | *Spinimuricea klavereni* | *Spinimuricea klavereni* |
| | | *Swiftia pallida* | DD | *Callistephanus pallida* | *Swiftia pallida* |
| Antipatharia | | | | | |
| | Antipathidae | *Antipathes dichotoma* | LC | *Antipathes dichotoma* | *Antipathes dichotoma* |
| | Leiopathidae | *Leiopathes glaberrima* | EN | *Leiopathes glaberrima* | *Leiopathes glaberrima* |
| | Myriopathidae | *Antipathella subpinnata* | LC | *Antipathella subpinnata* | *Antipathella subpinnata* |
| | Schizopathidae | *Parantipathes larix* | LC | *Parantipathes larix* | *Parantipathes larix* |
| Ceriantharia | | | | | |
| | Arachnactidae | *Arachnanthus oligopodus* | DD | *Arachnanthus oligopodus* | *Arachnanthus oligopodus* |
| Coleoptera | | | | | |
| | Alexiidae | *Sphaerosoma latitarse* | DD | *Sphaerosoma latitarse* | *Sphaerosoma latitarse* |
| | Anthribidae | *Anthribus fasciatus* | LC | *Anthribus fasciatus* | *Anthribus fasciatus* |
| | | *Anthribus nebulosus* | LC | *Anthribus nebulosus* | *Anthribus nebulosus* |
| | | *Anthribus scapularis* | DD | *Anthribus scapularis* | *Anthribus scapularis* |
| | | *Araecerodes grenieri* | LC | *Araecerodes grenieri* | *Araecerodes grenieri* |
| | | *Choragus aureolineatus* | DD | *Choragus aureolineatus* | *Choragus aurolineatus* |
| | | *Enedreytes hilaris* | LC | *Enedreytes hilaris* | *Enedreytes hilaris* |
| | | *Enedreytes sepicola* | LC | *Enedreytes sepicola* | *Pseudeuparius sepicola* |
| | | *Eusphyrus vasconicus* | DD | *Eusphyrus vasconicus* | *Eusphyrus vasconicus* |
| | | *Platystomos albinus* | LC | *Platystomos albinus* | *Platystomos albinus* |
| | | *Trigonorhinus areolatus* | DD | *Trigonorhinus areolatus* | *Trigonorhinus areolatus* |
| | Bostrichidae | *Amphicerus bimaculatus* | LC | *Amphicerus bimaculatus* | *Amphicerus bimaculatus* |
| | | *Xylomedes coronata* | DD | *Xylomedes coronata* | *Xylomedes coronata* |
| | Bothrideridae | *Teredus opacus* | VU | *Teredus opacus* | *Teredus opacus* |
| | Buprestidae | *Agrilus (Anambus) cyanescens ssp. italicus* | LC | *Agrilus cyanescens italicus* | *Agrilus cyanescens* |
| | | *Agrilus (Anambus) graecus* | DD | *Agrilus graecus* | *Agrilus graecus* |
| | | *Agrilus (Anambus) relegatus ssp. alexeevi* | LC | *Agrilus relegatus alexeevi* | *Agrilus relegatus subsp. alexeevi* |

| | | | | | |
|---|---|---|---|---|---|
| | | *Anthaxia (Anthaxia) candens* | VU | *Anthaxia candens* | *Anthaxia candens* |
| | | *Anthaxia (Anthaxia) chevrieri* | LC | *Anthaxia chevrieri* | *Anthaxia chevrieri* |
| | | *Anthaxia (Anthaxia) midas ssp. oberthuri* | EN | *Anthaxia midas oberthuri* | *Anthaxia midas subsp. oberthuri* |
| | | *Anthaxia (Anthaxia) spinolae* | LC | *Anthaxia spinolae* | *Anthaxia spinolae* |
| | | *Anthaxia (Anthaxia) suzannae* | LC | *Anthaxia suzannae* | *Anthaxia suzannae* |
| | | *Anthaxia (Haplanthaxia) flaviae* | LC | *Anthaxia flaviae* | *Anthaxia flaviae* |
| | | *Anthaxia (Melanthaxia) liae* | LC | *Anthaxia liae* | *Anthaxia liae* |
| | | *Anthaxia (Melanthaxia) rugicollis* | VU | *Anthaxia rugicollis* | *Anthaxia rugicollis* |
| | | *Coraebus fasciatus* | LC | *Coraebus fasciatus* | *Coraebus fasciatus* |
| | | *Lamprodila (Lamprodila) solieri* | CR | *Lamprodila solieri* | *Lamprodila solieri* |
| | | *Meliboeus (Meliboeus) fulgidicollis* | LC | *Meliboeus fulgidicollis* | *Meliboeus fulgidicollis* |
| | | *Perotis lugubris ssp. meridionalis* | NT | *Perotis lugubris meridionalis* | *Perotis lugubris subsp. meridionalis* |
| | | *Trachypteris picta ssp. decostigma* | LC | *Trachypteris picta decostigma* | *Trachypteris picta subsp. decostigma* |
| | Cerambycidae | *Acanthocinus henschi ssp. aetnensis* | CR | *Acanthocinus henschi aetnensis* | *Acanthocinus henschi subsp. aetnensis* |
| | | *Aegosoma scabricornis* | LC | *Aegosoma scabricornis* | *Aegosoma scabricorne* |
| | | *Anaglyptus zappii* | VU | *Anaglyptus zappii* | *Anaglyptus zappii* |
| | | *Cerambyx scopolii ssp. siculus* | NT | *Cerambyx scopolii siculus* | *Cerambyx scopolii subsp. siculus* |
| | | *Chlorophorus glaucus* | EN | *Chlorophorus glaucus* | *Chlorophorus glabromaculatus glaucus* |
| | | *Cornumutila lineata* | CR | *Cornumutila lineata* | *Cornumutila lineata* |
| | | *Etorofus pubescens* | NT | *Etorofus pubescens* | *Etorofus pubescens* |
| | | *Leioderes kollari ssp. jacopoi* | CR | *Leioderes kollari jacopoi* | *Leioderes kollari subsp. jacopoi* |
| | | *Liodерina linearis* | CR | *Liodерina linearis* | *Liodерina linearis* |
| | | *Morimus funereus* | VU | *Morimus funereus* | *Morimus asper funereus* |
| | | *Pogonocherus ovatoides* | CR | *Pogonocherus ovatoides* | *Pogonocherus ovatoides* |
| | | *Purpuricenus (Purpuricenus) apiceniger* | CR | *Purpuricenus apiceniger* | *Purpuricenus apiceniger* |
| | | *Rusticoclytus pantherinus* | EN | *Rusticoclytus pantherinus* | *Xylotrechus pantherinus* |
| | | *Rusticoclytus rusticus* | LC | *Rusticoclytus rusticus* | *Xylotrechus rusticus* |
| | | *Stenurella sennii* | DD | *Stenurella sennii* | *Stenurella melanura* |
| | | *Stictoleptura cordigera ssp. illyrica* | NT | *Stictoleptura cordigera illyrica* | *Stictoleptura cordigera subsp. illyrica* |
| | | *Trichoferus pallidus* | EN | *Trichoferus pallidus* | *Trichoferus pallidus* |

| | | | | | |
|---|---|---|---|---|---|
| | Cleridae | *Dermestoides sanguinicollis* | EN | *Dermestoides sanguinicollis* | *Dermestoides sanguinicollis* |
| | | *Opilo orocastaneus* | EN | *Opilo orocastaneus* | *Opilo orocastaneus* |
| | | *Teloclerus compressicornis* | VU | *Teloclerus compressicornis* | *Teloclerus compressicornis* |
| | Corylophidae | *Arthrolips nana* | DD | *Arthrolips nana* | *Arthrolips nana* |
| | | *Arthrolips obscura* | DD | *Arthrolips obscura* | *Arthrolips obscura* |
| | | *Clypastrea brunnea* | DD | *Clypastrea brunnea* | *Clypastrea brunnea* |
| | | *Clypastrea lata* | DD | *Clypastrea lata* | *Clypastrea lata* |
| | | *Clypastrea pusilla* | DD | *Clypastrea pusilla* | *Clypastrea pusilla* |
| | | *Clypastrea reitteri* | DD | *Clypastrea reitteri* | *Clypastrea reitteri* |
| | | *Orthoperus corticalis* | DD | *Orthoperus corticalis* | *Orthoperus corticalis* |
| | | *Orthoperus rogeri* | DD | *Orthoperus rogeri* | *Orthoperus rogeri* |
| | Cryptophagidae | *Atomaria (Agathengis) vespertina* | DD | *Atomaria vespertina* | *Atomaria vespertina* |
| | | *Caenoscelis angelinii* | CR | *Caenoscelis angelinii* | *Caenoscelis angelinii* |
| | | *Cryptophagus cylindrellus* | DD | *Cryptophagus cylindrellus* | *Cryptophagus cylindrellus* |
| | | *Cryptophagus denticulatus* | LC | *Cryptophagus denticulatus* | *Cryptophagus denticulatus* |
| | | *Cryptophagus parallelus* | DD | *Cryptophagus parallelus* | *Cryptophagus parallelus* |
| | | *Cryptophagus puncticollis* | DD | *Cryptophagus puncticollis* | *Cryptophagus puncticollis* |
| | | *Cryptophagus punctipennis* | LC | *Cryptophagus punctipennis* | *Cryptophagus punctipennis* |
| | | *Cryptophagus reflexus* | LC | *Cryptophagus reflexus* | *Cryptophagus reflexus* |
| | | *Cryptophagus schmidtii* | DD | *Cryptophagus schmidtii* | *Cryptophagus schmidtii* |
| | | *Cryptophagus schroetteri* | DD | *Cryptophagus schroetteri* | *Cryptophagus schroetteri* |
| | | *Cryptophagus uncinatus* | LC | *Cryptophagus uncinatus* | *Cryptophagus uncinatus* |
| | | *Micrambe pilosula* | DD | *Micrambe pilosula* | *Micrambe pilosula* |
| | Cucujidae | *Cucujus tulliae* | EN | *Cucujus tulliae* | *Cucujus tulliae* |
| | Curculionidae | *Acalles lemur ssp. cisalpinus* | LC | *Acalles lemur cisalpinus* | *Acalles lemur subsp. cisalpinus* |
| | | *Acalles sardiniaensis* | NT | *Acalles sardiniaensis* | *Acalles sardiniaensis* |
| | | *Acalles temperei* | NT | *Acalles temperei* | *Acalles temperei* |
| | | *Acallorneuma ingoi* | NT | *Acallorneuma ingoi* | *Acallorneuma ingoi* |
| | | *Acallorneuma montisalbi* | VU | *Acallorneuma montisalbi* | *Acallorneuma montisalbi* |
| | | *Acallorneuma sardeanense* | VU | *Acallorneuma sardeanense* | *Acallorneuma sardeanense* |
| | | *Anisandrus dispar* | LC | *Anisandrus dispar* | *Anisandrus dispar* |
| | | *Cryphalus asperatus* | LC | *Cryphalus asperatus* | *Cryphalus asperatus* |
| | | *Cryphalus numidicus* | LC | *Cryphalus numidicus* | *Cryphalus numidicus* |
| | | *Dichromacalles rolletii* | NT | *Dichromacalles rolletii* | *Dichromacalles rolletii* |
| | | *Dryocoetes italus* | DD | *Dryocoetes italus* | *Dryocoetes italus* |
| | | *Echinodera ibleiensis* | NT | *Echinodera ibleiensis* | *Echinodera ibleiensis* |

| | | | | | |
|---|---|---|---|---|---|
| | | *Echinodera kostenbaderi* | NT | *Echinodera kostenbaderi* | *Echinodera kostenbaderi* |
| | | *Echinodera nebrodiensis* | NT | *Echinodera nebrodiensis* | *Echinodera nebrodiensis* |
| | | *Echinodera peragalloi* | LC | *Echinodera peragalloi* | *Echinodera peragalloi* |
| | | *Echinodera settefratelliensis* | NT | *Echinodera settefratelliensis* | *Echinodera settefratelliensis* |
| | | *Echinodera siciliensis* | NT | *Echinodera siciliensis* | *Echinodera siciliensis* |
| | | *Ernoporicus fagi* | LC | *Ernoporicus fagi* | *Ernoporicus fagi* |
| | | *Gasterocercus depressirostris* | NT | *Gasterocercus depressirostris* | *Gasterocercus depressirostris* |
| | | *Gnathotrichus materiarius* | VU | *Gnathotrichus materiarius* | *Gnathotrichus materiarius* |
| | | *Hylastes gergeri* | DD | *Hylastes gergeri* | *Hylastes gergeri* |
| | | *Hylesinus toranio* | LC | *Hylesinus toranio* | *Hylesinus toranio* |
| | | *Hylesinus varius* | LC | *Hylesinus varius* | *Hylesinus varius* |
| | | *Hylobius (Callirus) abietis* | LC | *Hylobius abietis* | *Hylobius abietis* |
| | | *Hylobius (Callirus) pinastri* | LC | *Hylobius pinastri* | *Hylobius pinastri* |
| | | *Hylobius (Callirus) transversovittatus* | LC | *Hylobius transversovittatus* | *Hylobius transversovittatus* |
| | | *Hylobius (Hylobius) excavatus* | LC | *Hylobius excavatus* | *Hylobius excavatus* |
| | | *Kyklioacalles (Kyklioacalles) barbarus* | NT | *Kyklioacalles barbarus* | *Kyklioacalles barbarus* |
| | | *Kyklioacalles (Kyklioacalles) characivorus* | LC | *Kyklioacalles characivorus* | *Kyklioacalles characivorus* |
| | | *Kyklioacalles (Kyklioacalles) fausti* | LC | *Kyklioacalles fausti* | *Kyklioacalles fausti* |
| | | *Kyklioacalles (Kyklioacalles) provincialis* | VU | *Kyklioacalles provincialis* | *Kyklioacalles provincialis* |
| | | *Kyklioacalles (Kyklioacalles) punctaticollis* | LC | *Kyklioacalles punctaticollis* | *Kyklioacalles punctaticollis* |
| | | *Kyklioacalles (Kyklioacalles) punctaticollis ssp. meteoricus* | VU | *Kyklioacalles punctaticollis meteoricus* | *Kyklioacalles punctaticollis subsp. meteoricus* |
| | | *Kyklioacalles (Kyklioacalles) saccoi* | NT | *Kyklioacalles saccoi* | *Kyklioacalles saccoi* |
| | | *Kyklioacalles (Kyklioacalles) solarii* | LC | *Kyklioacalles solarii* | *Kyklioacalles solarii* |
| | | *Kyklioacalles (Kyklioacalles) teter* | NT | *Kyklioacalles teter* | *Kyklioacalles teter* |
| | | *Kyklioacalles (Palaeoacalles) navieresi* | LC | *Kyklioacalles navieresi* | *Kyklioacalles navieresi* |
| | | *Kyklioacalles (Palaeoacalles) roboris* | LC | *Kyklioacalles roboris* | *Kyklioacalles roboris* |
| | | *Lymantor coryli* | VU | *Lymantor coryli* | *Lymantor coryli* |
| | | *Melicius cylindrus* | LC | *Melicius cylindrus* | *Melicius cylindrus* |
| | | *Melicius gracilis* | LC | *Melicius gracilis* | *Melicius gracilis* |

| | | | | | |
|---|---|---|---|---|---|
| | | *Onyxacalles croaticus* | NT | *Onyxacalles croaticus* | *Onyxacalles croaticus* |
| | | *Onyxacalles henoni* | NT | *Onyxacalles henoni* | *Onyxacalles henoni* |
| | | *Onyxacalles luigionii* | LC | *Onyxacalles luigionii* | *Onyxacalles luigionii* |
| | | *Onyxacalles pyrenaeus* | LC | *Onyxacalles pyrenaeus* | *Onyxacalles pyrenaeus* |
| | | *Orthotomicus mannsfeldi* | LC | *Orthotomicus mannsfeldi* | *Orthotomicus mannsfeldi* |
| | | *Phloeotribus cristatus* | LC | *Phloeotribus cristatus* | *Phloeotribus cristatus* |
| | | *Phloeotribus perfoliatus* | LC | *Phloeotribus perfoliatus* | *Phloeotribus perfoliatus* |
| | | *Phloeotribus pubifrons* | LC | *Phloeotribus pubifrons* | *Phloeotribus pubifrons* |
| | | *Phloeotribus rhododactylus* | LC | *Phloeotribus rhododactylus* | *Phloeotribus rhododactylus* |
| | | *Phloeotribus spinulosus* | LC | *Phloeotribus spinulosus* | *Phloeotribus spinulosus* |
| | | *Pityokteines spinidens* | LC | *Pityokteines spinidens* | *Pityokteines spinidens* |
| | | *Pityophthorus carniolicus* | LC | *Pityophthorus carniolicus* | *Pityophthorus carniolicus* |
| | | *Scolytus ratzeburgii* | DD | *Scolytus ratzeburgii* | *Scolytus ratzeburgii* |
| | | *Stenoscelis (Stenoscelis) submuricata* | LC | *Stenoscelis submuricata* | *Stenoscelis submuricata* |
| | | *Thamnurgus kaltenbachii* | LC | *Thamnurgus kaltenbachii* | *Thamnurgus kaltenbachii* |
| | | *Trachodes hispidus* | LC | *Trachodes hispidus* | *Trachodes hispidus* |
| | | *Treptoplatypus oxyurus* | LC | *Treptoplatypus oxyurus* | *Treptoplatypus oxyurus* |
| | | *Triotemnus ulianai* | DD | *Triotemnus ulianai* | *Triotemnus ulianai* |
| | | *Trypodendron domesticum* | LC | *Trypodendron domesticum* | *Trypodendron domesticum* |
| | | *Trypodendron lineatum* | LC | *Trypodendron lineatum* | *Trypodendron lineatum* |
| | | *Trypodendron signatum* | LC | *Trypodendron signatum* | *Trypodendron signatum* |
| | | *Trypophloeus binodulus* | LC | *Trypophloeus binodulus* | *Trypophloeus binodulus* |
| | | *Xyleborinus saxesenii* | LC | *Xyleborinus saxesenii* | *Xyleborinus saxesenii* |
| | | *Xyleborus pfeili* | VU | *Xyleborus pfeili* | *Xyleborus pfeili* |
| | Dermestidae | *Globicornis (Globicornis) luckowi* | NT | *Globicornis luckowi* | *Globicornis luckowi* |
| | Elateridae | *Ampedus melonii* | VU | *Ampedus melonii* | *Ampedus melonii* |
| | | *Cardiophorus aetnensis* | DD | *Cardiophorus aetnensis* | *Cardiophorus aetnensis* |
| | | *Drapetes mordelloides* | LC | *Drapetes mordelloides* | *Drapetes mordelloides* |
| | | *Haterumelater fulvago* | EN | *Haterumelater fulvago* | *Haterumelater fulvago* |
| | | *Megathous valtopinensis* | EN | *Megathous valtopinensis* | *Megathous valtopinensis* |
| | Endomychidae | *Lycoperdina succinta* | NT | *Lycoperdina succinta* | *Lycoperdina succinta* |
| | Erotylidae | *Aulacochilus violaceus* | VU | *Aulacochilus violaceus* | *Aulacochilus violaceus* |
| | | *Cryptophilus integer* | LC | *Cryptophilus integer* | *Cryptophilus integer* |

| | | | | | |
|---|---|---|---|---|---|
| | | *Triplax andreinii* | DD | *Triplax andreinii* | *Triplax andreinii* |
| | | *Triplax lacordairii* | NT | *Triplax lacordairii* | *Triplax lacordairii* |
| | Eucnemidae | *Hylis cariniceps* | NT | *Hylis cariniceps* | *Hylis cariniceps* |
| | | *Hylis olexai* | NT | *Hylis olexai* | *Hylis olexai* |
| | | *Hylis procerulus* | DD | *Hylis procerulus* | *Hylis procerulus* |
| | | *Hylis simonae* | NT | *Hylis simonae* | *Hylis simonae* |
| | | *Microrhagus emyi* | VU | *Microrhagus emyi* | *Microrhagus emyi* |
| | | *Microrhagus hummleri* | CR | *Microrhagus hummleri* | *Microrhagus hummleri* |
| | | *Microrhagus lepidus* | NT | *Microrhagus lepidus* | *Microrhagus lepidus* |
| | | *Microrhagus pygmaeus* | NT | *Microrhagus pygmaeus* | *Microrhagus pygmaeus* |
| | | *Rhacopus sahlbergi* | DD | *Rhacopus sahlbergi* | *Rhacopus sahlbergi* |
| | | *Xylophilus corticalis* | NT | *Xylophilus corticalis* | *Xylophilus corticalis* |
| | | *Xylophilus testaceus* | EN | *Xylophilus testaceus* | *Xylophilus testaceus* |
| | Histeridae | *Abraeus globosus* | LC | *Abraeus globosus* | *Abraeus globosus* |
| | | *Epierus italicus* | LC | *Epierus italicus* | *Epierus italicus* |
| | | *Eubrachium pusillum* | LC | *Eubrachium pusillum* | *Eubrachium pusillum* |
| | | *Teretrius (Teretrius) picipes* | LC | *Teretrius picipes* | *Teretrius picipes* |
| | Latridiidae | *Cartodere nodifer* | LC | *Cartodere nodifer* | *Cartodere nodifer* |
| | | *Dienerella polyhymnia* | LC | *Dienerella polyhymnia* | *Dienerella polyhymnia* |
| | | *Dienerella vincenti* | DD | *Dienerella vincenti* | *Dienerella vincenti* |
| | | *Latridius amplus* | DD | *Latridius amplus* | *Latridius amplus* |
| | | *Latridius assimilis* | DD | *Latridius assimilis* | *Latridius assimilis* |
| | | *Stephostethus sinuaticollis* | DD | *Stephostethus sinuaticollis* | *Stephostethus sinuaticollis* |
| | Lucanidae | *Aesalus scarabaeoides ssp. siculus* | CR | *Aesalus scarabaeoides siculus* | *Aesalus scarabaeoides subsp. siculus* |
| | | *Lucanus tetraodon ssp. sicilianus* | NT | *Lucanus tetraodon sicilianus* | *Lucanus tetraodon subsp. sicilianus* |
| | Lycidae | *Lopherus rubens* | NT | *Lopherus rubens* | *Lopherus rubens* |
| | Lymexylidae | *Elateroides dermestoides* | NT | *Elateroides dermestoides* | *Elateroides dermestoides* |
| | Melandryidae | *Abdera (Abdera) bifasciata* | LC | *Abdera bifasciata* | *Abdera bifasciata* |
| | | *Dolotarsus lividus* | NT | *Dolotarsus lividus* | *Dolotarsus lividus* |
| | | *Phloiotrya (Phloiotrya) tenuis* | NT | *Phloiotrya tenuis* | *Phloiotrya tenuis* |
| | | *Zilora obscura* | VU | *Zilora obscura* | *Zilora obscura* |
| | Melyridae | *Aplocnemus (Aplocnemus) etruscus* | NT | *Aplocnemus etruscus* | *Aplocnemus etruscus* |
| | | *Aplocnemus (Aplocnemus) quercicola* | VU | *Aplocnemus quercicola* | *Aplocnemus quercicola* |
| | | *Aplocnemus (Diplambe) januaventi* | LC | *Aplocnemus januaventi* | *Aplocnemus januaventi* |
| | | *Dasytes (Dasytes) thoracicus ssp. lucanus* | VU | *Dasytes thoracicus lucanus* | *Dasytes thoracicus subsp. lucanus* |
| | | *Dasytes (Mesodasytes) aeneiventris* | LC | *Dasytes aeneiventris* | *Dasytes aeneiventris* |

| | | | | | |
|---|---|---|---|---|---|
| | | *Dasytes (Mesodasytes) croceipes* | LC | *Dasytes croceipes* | *Dasytes croceipes* |
| | | *Dasytes (Mesodasytes) nigrocyaneus* | LC | *Dasytes nigrocyaneus* | *Dasytes nigrocyaneus* |
| | | *Dasytes (Mesodasytes) virens* | LC | *Dasytes virens* | *Dasytes virens* |
| | | *Sphinginus lobatus ssp. apicalis* | LC | *Sphinginus lobatus apicalis* | *Sphinginus lobatus subsp. apicalis* |
| | Monotomidae | *Monotoma (Monotoma) quadricollis* | DD | *Monotoma quadricollis* | *Monotoma quadricollis* |
| | | *Rhizophagus (Cyanostolus) aeneus* | DD | *Rhizophagus aeneus* | *Rhizophagus aeneus* |
| | Mordellidae | *Mordellochroa milleri* | CR | *Mordellochroa milleri* | *Mordellochroa milleri* |
| | | *Pelecotoma fennica* | DD | *Pelecotoma fennica* | *Pelecotoma fennica* |
| | Mycetophagidae | *Esarcus (Entoxylon) abeillei* | NT | *Esarcus abeillei* | *Esarcus abeillei* |
| | | *Mycetophagus (Ulolendus) salicis* | NT | *Mycetophagus salicis* | *Mycetophagus salicis* |
| | | *Typhaea angusta* | DD | *Typhaea angusta* | *Typhaea angusta* |
| | Nitidulidae | *Epuraea unicolor* | LC | *Epuraea unicolor* | *Epuraea unicolor* |
| | | *Pityophagus quercus* | EN | *Pityophagus quercus* | *Pityophagus quercus* |
| | Oedemeridae | *Ischnomera caerulea* | LC | *Ischnomera caerulea* | *Ischnomera caerulea* |
| | | *Stenostoma cossyrense* | NT | *Stenostoma cossyrense* | *Stenostoma cossyrense* |
| | | *Stenostoma rostratum* | NT | *Stenostoma rostratum* | *Namibiana rostrata* |
| | Phloiophilidae | *Phloiophilus edvardsii* | DD | *Phloiophilus edvardsii* | *Phloiophilus edvardsii* |
| | Ptiliidae | *Actidium reitteri* | DD | *Actidium reitteri* | *Actidium reitteri* |
| | | *Ptenidium (Matthewsium) ponteleccianum* | NT | *Ptenidium ponteleccianum* | *Ptenidium ponteleccianum* |
| | | *Ptiliola brevicollis* | DD | *Ptiliola brevicollis* | *Ptiliola brevicollis* |
| | | *Ptinella mekula* | DD | *Ptinella mekula* | *Ptinella mekula* |
| | Ptinidae | *Anobium inexpectatum* | NT | *Anobium inexpectatum* | *Anobium inexpectatum* |
| | | *Ernobius gigas* | EN | *Ernobius gigas* | *Ernobius gigas* |
| | | *Gastrallus kocheri* | VU | *Gastrallus kocheri* | *Gastrallus kocheri* |
| | | *Gastrallus mauritanicus* | VU | *Gastrallus mauritanicus* | *Gastrallus mauritanicus* |
| | | *Hemicoelus canaliculatus* | LC | *Hemicoelus canaliculatus* | *Hemicoelus canaliculatus* |
| | | *Hyperisus declive* | EN | *Hyperisus declive* | *Hyperisus declive* |
| | | *Hyperisus plumbeum* | LC | *Hyperisus plumbeum* | *Hyperisus plumbeum* |
| | | *Metholcus phoenicius* | LC | *Metholcus phoenicius* | *Metholcus phoenicius* |
| | | *Priartobium serrifunis* | EN | *Priartobium serrifunis* | *Priartobium serrifunis* |
| | | *Xyletinus (Xyletinus) balcanicus* | VU | *Xyletinus balcanicus* | *Xyletinus balcanicus* |
| | Salpingidae | *Rabdocerus foveolatus* | LC | *Rabdocerus foveolatus* | *Rabdocerus foveolatus* |
| | | *Rabdocerus gabrieli* | NT | *Rabdocerus gabrieli* | *Rabdocerus gabrieli* |
| | | *Salpingus aeneus* | LC | *Salpingus aeneus* | *Salpingus aeneus* |
| | | *Salpingus planirostris* | LC | *Salpingus planirostris* | *Salpingus planirostris* |

| | | | | | |
|---|---|---|---|---|---|
| | | *Salpingus ruficollis* | NT | *Salpingus ruficollis* | *Salpingus ruficollis* |
| | | *Salpingus tapirus* | NT | *Salpingus tapirus* | *Salpingus tapirus* |
| | | *Sphaeriestes (Sphaeriestes) aeratus* | NT | *Sphaeriestes aeratus* | *Sphaeriestes aeratus* |
| | | *Sphaeriestes (Sphaeriestes) bimaculatus* | VU | *Sphaeriestes bimaculatus* | *Sphaeriestes bimaculatus* |
| | | *Sphaeriestes (Sphaeriestes) castaneus* | NT | *Sphaeriestes castaneus* | *Sphaeriestes castaneus* |
| | | *Sphaeriestes (Sphaeriestes) reyi* | NT | *Sphaeriestes reyi* | *Sphaeriestes reyi* |
| | | *Sphaeriestes (Sphaeriestes) stockmanni* | NT | *Sphaeriestes stockmanni* | *Sphaeriestes stockmanni* |
| | Scarabaeidae | *Aethiessa squamosa* | NT | *Aethiessa squamosa* | *Aethiessa squamosa* |
| | | *Calicnemis latreillii* | VU | *Calicnemis latreillii* | *Calicnemis latreillii* |
| | | *Osmoderma italicum* | EN | *Osmoderma italicum* | *Osmoderma italicum* |
| | | *Protaetia affinis* | LC | *Protaetia affinis* | *Protaetia affinis* |
| | | *Protaetia angustata* | DD | *Protaetia angustata* | *Protaetia angustata* |
| | | *Protaetia cuprea* | LC | *Protaetia cuprea* | *Protaetia cuprea* |
| | | *Protaetia cuprea ssp. hypocrita* | LC | *Protaetia cuprea hypocrita* | *Protaetia cuprea subsp. hypocrita* |
| | | *Protaetia fieberi* | VU | *Protaetia fieberi* | *Protaetia fieberi* |
| | | *Protaetia lugubris* | VU | *Protaetia lugubris* | *Protaetia lugubris* |
| | | *Protaetia mirifica* | CR | *Protaetia mirifica* | *Protaetia mirifica* |
| | | *Protaetia oblonga* | NT | *Protaetia oblonga* | *Protaetia oblonga* |
| | | *Protaetia opaca* | LC | *Protaetia opaca* | *Protaetia opaca* |
| | | *Protaetia sardea* | VU | *Protaetia sardea* | *Protaetia sardea* |
| | | *Protaetia speciosissima* | LC | *Protaetia speciosissima* | *Protaetia speciosissima* |
| | | *Protaetia squamosa* | VU | *Protaetia squamosa* | *Protaetia squamosa* |
| | | *Trichius gallicus* | LC | *Trichius gallicus* | *Trichius gallicus* |
| | | *Trichius gallicus ssp. zonatus* | CR | *Trichius gallicus zonatus* | *Trichius gallicus subsp. zonatus* |
| | Silvanidae | *Uleiota planatus* | LC | *Uleiota planatus* | *Uleiota planatus* |
| | Sphindidae | *Aspidiphorus lareyiniei* | NT | *Aspidiphorus lareyiniei* | *Aspidiphorus lareyiniei* |
| | | *Aspidiphorus orbiculatus* | LC | *Aspidiphorus orbiculatus* | *Aspidiphorus orbiculatus* |
| | | *Odontosphindus grandis* | VU | *Odontosphindus grandis* | *Odontosphindus grandis* |
| | Staphylinidae | *Amauronyx maerkeli* | NT | *Amauronyx maerkeli* | *Amauronyx maerkeli* |
| | | *Atheta liturata* | LC | *Atheta liturata* | *Atheta liturata* |
| | | *Atheta pallidicornis* | LC | *Atheta pallidicornis* | *Atheta pallidicornis* |
| | | *Atheta picipes* | LC | *Atheta picipes* | *Atheta picipes* |
| | | *Bolitochara humeralis* | NT | *Bolitochara humeralis* | *Bolitochara humeralis* |
| | | *Bolitochara tecta* | LC | *Bolitochara tecta* | *Bolitochara tecta* |
| | | *Caryoscapha limbata* | VU | *Caryoscapha limbata* | *Caryoscapha limbata* |
| | | *Dexiogyia corticina* | LC | *Dexiogyia corticina* | *Dexiogyia corticina* |
| | | *Dropephylla ammanni* | NT | *Dropephylla ammanni* | *Dropephylla ammanni* |

| | | | | | |
|---|---|---|---|---|---|
| | | *Dropephylla brevicornis* | NT | *Dropephylla brevicornis* | *Dropephylla brevicornis* |
| | | *Dropephylla devillei* | NT | *Dropephylla devillei* | *Dropephylla devillei* |
| | | *Dropephylla gracilicornis* | VU | *Dropephylla gracilicornis* | *Dropephylla gracilicornis* |
| | | *Dropephylla ioptera* | LC | *Dropephylla ioptera* | *Dropephylla ioptera* |
| | | *Dropephylla koltzei* | DD | *Dropephylla koltzei* | *Dropephylla koltzei* |
| | | *Dropephylla linearis* | VU | *Dropephylla linearis* | *Dropephylla linearis* |
| | | *Dropephylla perforata* | VU | *Dropephylla perforata* | *Dropephylla perforata* |
| | | *Dropephylla vilis* | NT | *Dropephylla vilis* | *Dropephylla vilis* |
| | | *Euryusa pipitzi* | CR | *Euryusa pipitzi* | *Euryusa pipitzi* |
| | | *Hypnogyra angularis* | LC | *Hypnogyra angularis* | *Hypnogyra angularis* |
| | | *Ischnoglossa prolixa* | NT | *Ischnoglossa prolixa* | *Ischnoglossa prolixa* |
| | | *Phloeopora corticalis* | LC | *Phloeopora corticalis* | *Phloeopora corticalis* |
| | | *Rugilus mixtus* | CR | *Rugilus mixtus* | *Rugilus mixtus* |
| | | *Sepedophilus aestivus* | NT | *Sepedophilus aestivus* | *Sepedophilus aestivus* |
| | Tenebrionidae | *Accanthopus velikensis* | LC | *Accanthopus velikensis* | *Accanthopus velikensis* |
| | | *Allecula suberina* | EN | *Allecula suberina* | *Allecula suberina* |
| | | *Corticeus bicolor* | LC | *Corticeus bicolor* | *Corticeus bicolor* |
| | | *Corticeus bicoloroides* | CR | *Corticeus bicoloroides* | *Corticeus bicoloroides* |
| | | *Corticeus fasciatus* | LC | *Corticeus fasciatus* | *Corticeus fasciatus* |
| | | *Corticeus linearis* | LC | *Corticeus linearis* | *Corticeus linearis* |
| | | *Corticeus pini* | LC | *Corticeus pini* | *Corticeus pini* |
| | | *Corticeus suberis* | DD | *Corticeus suberis* | *Corticeus suberis* |
| | | *Corticeus unicolor* | LC | *Corticeus unicolor* | *Corticeus unicolor* |
| | | *Corticeus versipellis* | DD | *Corticeus versipellis* | *Corticeus versipellis* |
| | | *Diaclina fagi* | DD | *Diaclina fagi* | *Diaclina fagi* |
| | | *Eledona agricola* | NT | *Eledona agricola* | *Eledona agricola* |
| | | *Eledonoprius serrifrons* | CR | *Eledonoprius serrifrons* | *Eledonoprius serrifrons* |
| | | *Mycetochara (Ernocharis) straussii* | CR | *Mycetochara straussii* | *Mycetochara straussii* |
| | | *Nalassus pastai* | CR | *Nalassus pastai* | *Nalassus pastai* |
| | | *Neomida haemorrhoidalis* | EN | *Neomida haemorrhoidalis* | *Neomida haemorrhoidalis* |
| | | *Odocnemis clypeatus* | NT | *Odocnemis clypeatus* | *Odocnemis clypeatus* |
| | | *Pelorinus ebeninus* | LC | *Pelorinus ebeninus* | *Pelorinus ebeninus* |
| | | *Platydema europaea* | CR | *Platydema europaea* | *Platydema europaea* |
| | | *Platydema violacea* | NT | *Platydema violacea* | *Platydema violacea* |
| | | *Prionychus fairmairii* | NT | *Prionychus fairmairii* | *Prionychus fairmairii* |
| | | *Scaphidema metallica* | LC | *Scaphidema metallica* | *Scaphidema metallica* |
| | Tetratomidae | *Tetratoma desmarestii* | EN | *Tetratoma desmarestii* | *Tetratoma desmarestii* |
| | | *Tetratoma tedaldi* | VU | *Tetratoma tedaldi* | *Tetratoma tedaldi* |
| | Throscidae | *Aulonothroscus brevicollis* | DD | *Aulonothroscus brevicollis* | *Aulonothroscus brevicollis* |

| | | | | | |
|---|---|---|---|---|---|
| | | *Trixagus algiricus* | DD | *Trixagus algiricus* | *Trixagus algiricus* |
| | | *Trixagus angelinii* | LC | *Trixagus angelinii* | *Trixagus angelinii* |
| | | *Trixagus asiaticus* | DD | *Trixagus asiaticus* | *Trixagus asiaticus* |
| | | *Trixagus atticus* | DD | *Trixagus atticus* | *Trixagus atticus* |
| | | *Trixagus carinifrons* | DD | *Trixagus carinifrons* | *Trixagus carinifrons* |
| | | *Trixagus dermestoides* | LC | *Trixagus dermestoides* | *Trixagus dermestoides* |
| | | *Trixagus duvalii* | DD | *Trixagus duvalii* | *Trixagus duvalii* |
| | | *Trixagus gracilis* | LC | *Trixagus gracilis* | *Trixagus gracilis* |
| | | *Trixagus leseigneuri* | DD | *Trixagus leseigneuri* | *Trixagus leseigneuri* |
| | | *Trixagus minutus* | DD | *Trixagus minutus* | *Trixagus minutus* |
| | | *Trixagus myebohmi* | NT | *Trixagus myebohmi* | *Trixagus myebohmi* |
| | | *Trixagus obtusus* | LC | *Trixagus obtusus* | *Trixagus obtusus* |
| | | *Trixagus rougeti* | DD | *Trixagus rougeti* | *Trixagus rougeti* |
| | Trogossitidae | *Nemozoma elongatum* | LC | *Nemozoma elongatum* | *Nemozoma elongatum* |
| | | *Ostoma ferruginea* | NT | *Ostoma ferruginea* | *Ostoma ferruginea* |
| | Zopheridae | *Endophloeus marcovichianus* | NT | *Endophloeus marcovichianus* | *Endophloeus marcovichianus* |
| | | *Langelandia anophtalma* | LC | *Langelandia anophtalma* | *Langelandia anophtalma* |
| | | *Langelandia montalbica* | CR | *Langelandia montalbica* | *Langelandia montalbica* |
| | | *Langelandia reitteri* | NT | *Langelandia reitteri* | *Langelandia reitteri* |
| | | *Langelandia vienensis* | DD | *Langelandia vienensis* | *Langelandia vienensis* |
| | | *Pycnomerus italicus* | EN | *Pycnomerus italicus* | *Pycnomerus italicus* |
| | | *Synchita fallax* | NT | *Synchita fallax* | *Synchita fallax* |
| | | *Synchita undata* | NT | *Synchita undata* | *Synchita undata* |
| | | *Synchita variegata* | LC | *Synchita variegata* | *Synchita variegata* |
| Hymenoptera | | | | | |
| | Andrenidae | *Andrena palumba* | EN | *Andrena palumba* | *Andrena palumba* |
| | | *Andrena aberrans* | DD | *Andrena aberrans* | *Andrena aberrans* |
| | | *Andrena binominata* | DD | *Andrena binominata* | *Andrena binominata* |
| | | *Andrena boyerella* | DD | *Andrena boyerella* | *Andrena boyerella* |
| | | *Andrena curvana* | DD | *Andrena curvana* | *Andrena curvana* |
| | | *Andrena diomedia* | DD | *Andrena diomedia* | *Andrena diomedia* |
| | | *Andrena incisa* | DD | *Andrena incisa* | *Andrena incisa* |
| | | *Andrena iohannescaroli* | DD | *Andrena iohannescaroli* | *Andrena iohannescaroli* |
| | | *Andrena panurgina* | DD | *Andrena panurgina* | *Andrena panurgina* |
| | | *Andrena suerinensis* | DD | *Andrena suerinensis* | *Andrena suerinensis* |
| | | *Panurgus corsicus* | DD | *Panurgus corsicus* | *Panurgus corsicus* |
| | Apidae | *Bombus konradini* | EN | *Bombus konradini* | *Bombus konradini* |
| | | *Nomada roberjeotiana* | VU | *Nomada roberjeotiana* | *Nomada roberjeotiana* |
| | | *Anthophora nigrovittata* | DD | *Anthophora nigrovittata* | *Anthophora balneorum* |
| | | *Bombus haematurus* | DD | *Bombus haematurus* | *Bombus haematurus* |
| | | *Bombus perezi* | DD | *Bombus perezi* | *Bombus perezi* |
| | | *Epeolus fasciatus* | DD | *Epeolus fasciatus* | *Epeolus fasciatus* |

| | | | | | |
|---|---|---|---|---|---|
| | | *Epeolus tarsalis* | DD | *Epeolus tarsalis* | *Epeolus tarsalis* |
| | | *Eucera albofasciata* | DD | *Eucera albofasciata* | *Eucera nigrita* |
| | | *Habropoda ezonata* | DD | *Habropoda ezonata* | *Habropoda ezonata* |
| | | *Nomada alpigena* | DD | *Nomada alpigena* | *Nomada alpigena* |
| | | *Nomada duplex* | DD | *Nomada duplex* | *Nomada duplex* |
| | | *Nomada pectoralis* | DD | *Nomada pectoralis* | *Nomada pectoralis* |
| | Colletidae | *Colletes collaris* | EN | *Colletes collaris* | *Colletes collaris* |
| | | *Colletes wolfi* | EN | *Colletes wolfi* | *Colletes wolfi* |
| | | *Colletes tuberculatus* | NT | *Colletes tuberculatus* | *Colletes tuberculatus* |
| | | *Colletes acutiformis* | DD | *Colletes acutiformis* | *Colletes acutiformis* |
| | | *Colletes foveolaris* | DD | *Colletes foveolaris* | *Colletes foveolaris* |
| | | *Colletes graeffei* | DD | *Colletes graeffei* | *Colletes graeffei* |
| | | *Lasioglossum algirum* | DD | *Lasioglossum algirum* | *Lasioglossum algirum* |
| | | *Sphecodes pinguiculus* | DD | *Sphecodes pinguiculus* | *Sphecodes pinguiculus* |
| | Megachilidae | *Pseudoanthidium eximium* | NT | *Pseudoanthidium eximium* | *Pseudoanthidium eximium* |
| | | *Chelostoma stefanii* | DD | *Chelostoma stefanii* | *Chelostoma stefanii* |
| | | *Heriades punctulifera* | DD | *Heriades punctulifera* | *Heriades punctulifera* |
| | | *Hoplitis occidentalis* | DD | *Hoplitis occidentalis* | *Hoplitis occidentalis* |
| | | *Hoplitis saxialis* | DD | *Hoplitis saxialis* | *Hoplitis saxialis* |
| | | *Megachile anatolica* | DD | *Megachile anatolica* | *Megachile anatolica* |
| | | *Megachile burdigalensis* | DD | *Megachile burdigalensis* | *Megachile burdigalensis* |
| | | *Osmia picena* | DD | *Osmia picena* | *Osmia picena* |
| | | *Rhodanthidium acuminatum* | DD | *Rhodanthidium acuminatum* | *Rhodanthidium acuminatum* |
| | | *Stenoheriades maroccana* | DD | *Stenoheriades maroccana* | *Stenoheriades maroccana* |
| | | *Trachusa interrupta* | DD | *Trachusa interrupta* | *Trachusa interrupta* |
| | Melittidae | *Macropis frivaldszkyi* | CR (PE) | *Macropis frivaldszkyi* | *Macropis frivaldszkyi* |
| | | *Dasypoda braccata* | CR | *Dasypoda braccata* | *Dasypoda braccata* |
| | | *Melitta tomentosa* | DD | *Melitta tomentosa* | *Melitta tomentosa* |
| Lepidoptera | | | | | |
| | Hesperiidae | *Spialia orbifera* | LC | *Spialia orbifera* | *Spialia orbifera* |
| | | *Sloperia proto* | LC | *Sloperia proto* | *Sloperia proto* |
| | | *Ochlodes sylvanus* | LC | *Ochlodes sylvanus* | *Ochlodes sylvanus* |
| | Papilionidae | *Parnassius sacerdos (phoebus)* | LC | *Parnassius sacerdos* | *Parnassius sacerdos* |
| | | *Zerynthia cassandra* | LC | *Zerynthia cassandra* | *Zerynthia cassandra* |
| | Pieridae | *Euchloe tagis (bellezina)* | NT | *Euchloe tagis* | *Euchloe tagis* |
| | | *Leptidea juvernica* | LC | *Leptidea juvernica* | *Leptidea juvernica* |
| | Lycaenidae | *Callophrys avis* | VU | *Callophrys avis* | *Callophrys avis* |
| | | *Zizeeria karsandra* | VU | *Zizeeria karsandra* | *Zizeeria karsandra* |
| | | *Azanus ubaldus* | DD | *Azanus ubaldus* | *Azanus ubaldus* |
| | | *Polyommatus celinus* | LC | *Polyommatus celinus* | *Polyommatus celinus* |
| | | *Polyommatus icarius* | LC | *Polyommatus icarius* | *Polyommatus icarus icarius* |

| | | | | | |
|---|---|---|---|---|---|
| | Nymphalidae | *Melitaea nevadensis* | LC | *Melitaea nevadensis* | *Mellicta parthenoides veletaensis* |
| | | *Melitaea ornata* | LC | *Melitaea ornata* | *Melitaea trivia* |
| | | *Erebia aethiopellus* | LC | *Erebia aethiopellus* | *Erebia mnestra* |
| | | *Erebia albergana* | LC | *Erebia albergana* | *Erebia albergana* |
| | | *Erebia dromus* | LC | *Erebia dromus* | *Erebia tyndarus* |
| | | *Coenonympha lyllus* | LC | *Coenonympha lyllus* | *Coenonympha lyllus* |
| | | *Lasiommata paramegaera* | LC | *Lasiommata paramegaera* | *Lasiommata paramegaera* |
| Pennatulacea | | | | | |
| | Kophobelemnidae | *Kophobelemnon stelliferum* | LC | *Kophobelemnon stelliferum* | *Kophobelemnon stelliferum* |
| Scleractinia | | | | | |
| | Caryophyllidae | *Phyllangia americana* | DD | *Phyllangia americana* | *Phyllangia americana* |
| | Oculinidae | *Oculina patagonica* | LC | *Oculina patagonica* | *Oculina patagonica* |
