## Supplementary Table 2 for "How well is Italian biodiversity represented in red lists and conservation legislation? Taxonomic biases, coverage gaps and the assessment-to-legislation bottleneck"

**Supplementary table 2:** Summary table of counts of assessed and non-assessed taxa in Italy for each of the assessments gathered in the study (ITA: Italian regional assessments, EU: European Red Lists, GLO: Global IUCN Red List, MED: Mediterranean assessments). When assessments were requested at iucn.org, all downloads are accurate to April 2025. The columns explicit the subtotals for endemics. Percentages for "Totals for major groups" refer to the percentage of each major group to the grand total of all taxa included in the study. In the columns differentiating assessed and non-assessed endemics, the percentages represent the proportion of taxa over the total number of endemics. Finally, for non-endemic taxa, the percentages refer to the proportion of taxa over the total of non-endemics.

| Major taxa | Taxa groups | Scope | Totals per major group | Total endemics | Total non endemics | Total assessed | Total non-assessed | Assessed endemics | Non assessed endemics | Assessed non-endemics | Non assessed non endemics |
|---|---|---|---|---|---|---|---|---|---|---|---|
| **Algae** | Charophyta | ITA | 35 (0.05%) | - | 35 (100%) | 0 (0%) | 35 (100%) | - | - | 0 (0%) | 35 (100%) |
| | | EU | | | | 12 (34%) | 23 (66%) | - | - | 12 (34%) | 23 (66%) |
| | | GLO | | | | 1 (3%) | 34 (97%) | - | - | 1 (3%) | 34 (97%) |
| | | MED | | | | 0 (0%) | 35 (100%) | - | - | 0 (0%) | 35 (100%) |
| | Chlorophyta | ITA | 71 (0.09%) | - | 71 (100%) | 0 (0%) | 71 (100%) | - | - | 0 (0%) | 71 (100%) |
| | | EU | | | | 0 (0%) | 71 (100%) | - | - | 0 (0%) | 71 (100%) |
| | | GLO | | | | 0 (0%) | 71 (100%) | - | - | 0 (0%) | 71 (100%) |
| | | MED | | | | 0 (0%) | 71 (100%) | - | - | 0 (0%) | 71 (100%) |
| | Ochrophyta | ITA | 130 (0.17%) | - | 130 (100%) | 0 (0%) | 130 (100%) | - | - | 0 (0%) | 130 (100%) |
| | | EU | | | | 0 (0%) | 130 (100%) | - | - | 0 (0%) | 130 (100%) |
| | | GLO | | | | 0 (0%) | 130 (100%) | - | - | 0 (0%) | 130 (100%) |
| | | MED | | | | 0 (0%) | 130 (100%) | - | - | 0 (0%) | 130 (100%) |
| | Rhodophyta | ITA | 293 (0.38%) | - | 293 (100%) | 0 (0%) | 293 (100%) | - | - | 0 (0%) | 293 (100%) |
| | | EU | | | | 0 (0%) | 293 (100%) | - | - | 0 (0%) | 293 (100%) |
| | | GLO | | | | 0 (0%) | 293 (100%) | - | - | 0 (0%) | 293 (100%) |
| | | MED | | | | 0 (0%) | 293 (100%) | - | - | 0 (0%) | 293 (100%) |
| **Insects** | Coleoptera | ITA | 13,148 (17%) | 2,608 (20%) | 10,540 (80%) | 1,749 (13%) | 11,399 (87%) | 136 (5%) | 2,472 (95%) | 1,613 (15%) | 8,927 (85%) |

| | | | | | | | | | | | |
|---|---|---|---|---|---|---|---|---|---|---|---|
| | | EU | | | | 399 (3%) | 12,749 (97%) | 31 (1%) | 2,577 (99%) | 368 (4%) | 10,172 (96%) |
| | | GLO | | | | 117 (1%) | 13,031 (99%) | 24 (1%) | 2,584 (99%) | 93 (1%) | 10,447 (99%) |
| | | MED | | | | 114 (1%) | 13,034 (99%) | 20 (1%) | 2,588 (99%) | 94 (1%) | 10,446 (99%) |
| | Diptera | ITA | 6,676 (8.7%) | 261 (4%) | 6,415 (96%) | 0 (0%) | 6,676 (100%) | 0 (0%) | 261 (100%) | 0 (0%) | 6,415 (100%) |
| | | EU | | | | 414 (6%) | 6,257 (94%) | 7 (3%) | 254 (97%) | 412 (6%) | 6,003 (94%) |
| | | GLO | | | | 95 (1%) | 6,579 (99%) | 7 (3%) | 254 (97%) | 90 (1%) | 6,325 (99%) |
| | | MED | | | | 11 (0%) | 6,664 (100%) | 3 (1%) | 258 (99%) | 9 (0%) | 6,406 (100%) |
| | Hymenoptera | ITA | 7,966 (10.37%) | 131 (2%) | 7,835 (98%) | 124 (2%) | 7,842 (98%) | 5 (4%) | 126 (96%) | 119 (2%) | 7,716 (98%) |
| | | EU | | | | 1,028 (13%) | 6,938 (87%) | 16 (12%) | 115 (88%) | 1012 (13%) | 6,823 (87%) |
| | | GLO | | | | 116 (1%) | 7,850 (99%) | 7 (5%) | 124 (95%) | 109 (1%) | 7,726 (99%) |
| | | MED | | | | 0 (0%) | 7,966 (100%) | 0 (0%) | 131 (100%) | 0 (0%) | 7,835 (100%) |
| | Lepidoptera | ITA | 5,168 (6.73%) | 160 (3%) | 5,008 (97%) | 268 (5%) | 4,900 (95%) | 15 (9%) | 145 (91%) | 253 (5%) | 4,755 (95%) |
| | | EU | | | | 249 (5%) | 4,919 (95%) | 10 (6%) | 150 (94%) | 239 (5%) | 4,769 (95%) |
| | | GLO | | | | 86 (2%) | 5,082 (98%) | 10 (6%) | 150 (94%) | 76 (2%) | 4,932 (98%) |
| | | MED | | | | 171 (3%) | 4,997 (97%) | 8 (5%) | 152 (95%) | 163 (3%) | 4,845 (97%) |
| | Mantodea | ITA | 10 (0.01%) | 1 (10%) | 9 (90%) | 0 (0%) | 10 (100%) | 0 (0%) | 1 (100%) | 0 (0%) | 9 (100%) |
| | | EU | | | | 3 (30%) | 7 (70%) | 1 (100%) | 0 | 2 (22%) | 7 (78%) |
| | | GLO | | | | 10 (100%) | 0 (0%) | 1 (100%) | 0 | 9 (100%) | 0 (0%) |
| | | MED | | | | 0 (0%) | 10 (100%) | 0 (0%) | 1 (100%) | 0 (0%) | 9 (100%) |
| | Odonata | ITA | 95 (0.1%) | 1 (1%) | 94 (99%) | 83 (87%) | 12 (13%) | 1 (100%) | 0 | 82 (87%) | 12 (13%) |
| | | EU | | | | 91 (96%) | 4 (4%) | 1 (100%) | 0 | 90 (96%) | 4 (4%) |
| | | GLO | | | | 83 (87%) | 12 (13%) | 1 (100%) | 0 | 82 (87%) | 12 (13%) |

| | | | | | | | | | | |
|---|---|---|---|---|---|---|---|---|---|---|
| | | MED | | | | 3 (3%) | 92 (97%) | 1 (100%) | 0 | 2 (2%) | 92 (98%) |
| | Orthoptera | ITA | 408 (0.5%) | 115 (28%) | 293 (72%) | 0 (0%) | 408 (100%) | 0 (0%) | 115 (100%) | 0 (0%) | 293 (100%) |
| | | EU | | | | 364 (89%) | 44 (11%) | 105 (91%) | 10 (9%) | 259 (88%) | 34 (12%) |
| | | GLO | | | | 190 (47%) | 218 (53%) | 89 (77%) | 26 (23%) | 101 (34%) | 192 (66%) |
| | | MED | | | | 101 (25%) | 307 (75%) | 70 (61%) | 45 (39%) | 31 (11%) | 262 (89%) |
| | Other insects | ITA | 4,425 (6%) | 453 (10%) | 3,972 (90%) | 0 (0%) | 4,425 (100%) | 0 (0%) | 453 (100%) | 0 (0%) | 3,972 (100%) |
| | | EU | | | | 0 (0%) | 4,425 (100%) | 0 (0%) | 453 (100%) | 0 (0%) | 3,972 (100%) |
| | | GLO | | | | 0 (0%) | 4,425 (100%) | 0 (0%) | 453 (100%) | 0 (0%) | 3,972 (100%) |
| | | MED | | | | 0 (0%) | 4,425 (100%) | 0 (0%) | 453 (100%) | 0 (0%) | 3,972 (100%) |
| **Lichens** | Ascolichen | ITA | 3,272 (4.26%) | - | 3,272 (100%) | 154 (5%) | 3,118 (95%) | - | - | 154 (5%) | 3,118 (95%) |
| | | EU | | | | 0 (0%) | 3,272 (100%) | - | - | 0 (0%) | 3,272 (100%) |
| | | GLO | | | | 7 (0%) | 3,265 (100%) | - | - | 7 (0%) | 3,265 (100%) |
| | | MED | | | | 0 (0%) | 3,272 (100%) | - | - | 0 (0%) | 3,272 (100%) |
| | Basidiolichen | ITA | 3 (0%) | - | 3 (100%) | 0 (0%) | 3 (100%) | - | - | 0 (0%) | 3 (100%) |
| | | EU | | | | 0 (0%) | 3 (100%) | - | - | 0 (0%) | 3 (100%) |
| | | GLO | | | | 0 (0%) | 3 (100%) | - | - | 0 (0%) | 3 (100%) |
| | | MED | | | | 0 (0%) | 3 (100%) | - | - | 0 (0%) | 3 (100%) |
| **Macrofungi** | Ascomycota | ITA | 289 (0.38%) | - | 289 (100%) | 2 (1%) | 287 (99%) | - | - | 2 (1%) | 287 (99%) |
| | | EU | | | | 0 (0%) | 289 (100%) | - | - | 0 (0%) | 289 (100%) |

| | | | | | | | | | | | |
|---|---|---|---|---|---|---|---|---|---|---|---|
| | | GLO | | | | 1 (0%) | 288 (100%) | - | - | 1 (0%) | 288 (100%) |
| | | MED | | | | 0 (0%) | 289 (100%) | - | - | 0 (0%) | 289 (100%) |
| | Basidiomycota | ITA | 4,168 (5.43%) | 6 (0%) | 4,162 (100%) | 19 (0%) | 4,149 (100%) | 0 (0%) | 6 (100%) | 19 (0%) | 4,143 (100%) |
| | | EU | | | | 2 (0%) | 4,166 (100%) | 0 (0%) | 6 (100%) | 2 (0%) | 4,160 (100%) |
| | | GLO | | | | 114 (3%) | 4,054 (97%) | 2 (33%) | 4 (67%) | 112 (3%) | 4,050 (97%) |
| | | MED | | | | 0 (0%) | 4,168 (100%) | 0 (0%) | 6 (100%) | 0 (0%) | 4,162 (100%) |
| **Molluscs** | Bivalvia | ITA | 370 (0.48%) | 10 (3%) | 360 (97%) | 0 (0%) | 370 (100%) | 0 (0%) | 10 (100%) | 0 (0%) | 360 (100%) |
| | | EU | | | | 25 (7%) | 345 (93%) | 0 (0%) | 10 (100%) | 25 (7%) | 335 (93%) |
| | | GLO | | | | 18 (5%) | 352 (95%) | 0 (0%) | 10 (100%) | 18 (5%) | 342 (95%) |
| | | MED | | | | 2 (1%) | 368 (99%) | 0 (0%) | 10 (100%) | 2 (1%) | 358 (99%) |
| | Cephalopoda | ITA | 39 (0.05%) | 4 (10%) | 35 (90%) | 0 (0%) | 39 (100%) | 0 (0%) | 4 (100%) | 0 (0%) | 35 (100%) |
| | | EU | | | | 0 (0%) | 39 (100%) | 0 (0%) | 4 (100%) | 0 (0%) | 35 (100%) |
| | | GLO | | | | 39 (100%) | 0 (0%) | 4 (100%) | 0 | 35 (100%) | 0 (0%) |
| | | MED | | | | 0 (0%) | 39 (100%) | 0 (0%) | 4 (100%) | 0 (0%) | 35 (100%) |
| | Gastropoda | ITA | 2,117 (2.76%) | 538 (25%) | 1,579 (75%) | 0 (0%) | 2,117 (100%) | 0 (0%) | 538 (100%) | 0 (0%) | 1,579 (100%) |
| | | EU | | | | 707 (33%) | 1,410 (67%) | 307 (57%) | 231 (43%) | 400 (25%) | 1,179 (75%) |
| | | GLO | | | | 611 (29%) | 1,506 (71%) | 299 (56%) | 239 (44%) | 312 (20%) | 1,267 (80%) |
| | | MED | | | | 130 (6%) | 1,987 (94%) | 80 (15%) | 458 (85%) | 50 (3%) | 1,529 (97%) |
| | Other molluscs | ITA | 80 (0.1%) | 2 (2%) | 78 (98%) | 0 (0%) | 80 (100%) | 0 (0%) | 2 (100%) | 0 (0%) | 78 (100%) |
| | | EU | | | | 0 (0%) | 80 (100%) | 0 (0%) | 2 (100%) | 0 (0%) | 78 (100%) |
| | | GLO | | | | 0 (0%) | 80 (100%) | 0 (0%) | 2 (100%) | 0 (0%) | 78 (100%) |
| | | MED | | | | 0 (0%) | 80 (100%) | 0 (0%) | 2 (100%) | 0 (0%) | 78 (100%) |

| | | | | | | | | | | | |
|---|---|---|---|---|---|---|---|---|---|---|---|
| **Non-vascular plants** | Andreaeopsida | ITA | 10 (0.01%) | - | 10 (100%) | 10 (100%) | 0 (0%) | - | - | 10 (100%) | 0 (0%) |
| | | EU | | | | 8 (80%) | 2 (20%) | - | - | 8 (80%) | 2 (20%) |
| | | GLO | | | | 2 (20%) | 8 (80%) | - | - | 2 (20%) | 8 (80%) |
| | | MED | | | | 0 (0%) | 10 (100%) | - | - | 0 (0%) | 10 (100%) |
| | Anthocerotopsida | ITA | 7 (0.01%) | - | 7 (100%) | 6 (86%) | 1 (14%) | - | - | 6 (86%) | 1 (14%) |
| | | EU | | | | 6 (86%) | 1 (14%) | - | - | 6 (86%) | 1 (14%) |
| | | GLO | | | | 0 (0%) | 7 (100%) | - | - | 0 (0%) | 7 (100%) |
| | | MED | | | | 0 (0%) | 7 (100%) | - | - | 0 (0%) | 7 (100%) |
| | Bryopsida | ITA | 898 (1.17%) | - | 898 (100%) | 889 (99%) | 9 (1%) | - | - | 889 (99%) | 9 (1%) |
| | | EU | | | | 840 (94%) | 58 (6%) | - | - | 840 (94%) | 58 (6%) |
| | | GLO | | | | 27 (3%) | 871 (97%) | - | - | 27 (3%) | 871 (97%) |
| | | MED | | | | 1 (0%) | 897 (100%) | - | - | 1 (0%) | 897 (100%) |
| | Marchantiophyta | ITA | 303 (0.4%) | - | 303 (100%) | 297 (98%) | 6 (2%) | - | - | 297 (98%) | 6 (2%) |
| | | EU | | | | 300 (99%) | 3 (1%) | - | - | 300 (99%) | 3 (1%) |
| | | GLO | | | | 6 (2%) | 297 (98%) | - | - | 6 (2%) | 297 (98%) |
| | | MED | | | | 0 (0%) | 303 (100%) | - | - | 0 (0%) | 303 (100%) |
| | Polytrichopsida | ITA | 21 (0.03%) | - | 21 (100%) | 21 (100%) | 0 (0%) | - | - | 21 (100%) | 0 (0%) |
| | | EU | | | | 20 (95%) | 1 (5%) | - | - | 20 (95%) | 1 (5%) |
| | | GLO | | | | 0 (0%) | 21 (100%) | - | - | 0 (0%) | 21 (100%) |
| | | MED | | | | 0 (0%) | 21 (100%) | - | - | 0 (0%) | 21 (100%) |
| | Sphagnopsida | ITA | 35 (0.05%) | - | 35 (100%) | 34 (97%) | 1 (3%) | - | - | 34 (97%) | 1 (3%) |
| | | EU | | | | 34 (97%) | 1 (3%) | - | - | 34 (97%) | 1 (3%) |
| | | GLO | | | | 0 (0%) | 35 (100%) | - | - | 0 (0%) | 35 (100%) |
| | | MED | | | | 0 (0%) | 35 (100%) | - | - | 0 (0%) | 35 (100%) |
| **Other invertebrates** | Anthozoa | ITA | 126 (0.16%) | 32 (25%) | 94 (75%) | 100 (79%) | 26 (21%) | 25 (78%) | 7 (22%) | 75 (80%) | 19 (20%) |
| | | EU | | | | 3 (2%) | 123 (98%) | 3 (9%) | 29 (91%) | 0 | 94 (100%) |

| | | | | | | | | | | | |
|---|---|---|---|---|---|---|---|---|---|---|---|
| | | GLO | | | | 21 (17%) | 105 (83%) | 12 (38%) | 20 (62%) | 9 (10%) | 85 (90%) |
| | | MED | | | | 87 (69%) | 39 (31%) | 22 (69%) | 10 (31%) | 65 (69%) | 29 (31%) |
| | Arachnida | ITA | 5,125 (6.67%) | 615 (12%) | 4,510 (88%) | 0 (0%) | 5,125 (100%) | 0 (0%) | 615 (100%) | 0 (0%) | 4,510 (100%) |
| | | EU | | | | 0 (0%) | 5,125 (100%) | 0 (0%) | 615 (100%) | 0 (0%) | 4,510 (100%) |
| | | GLO | | | | 2 (0.3%) | 5,123 (99%) | 1 (0%) | 614 (99%) | 1 (0.16%) | 4,509 (99%) |
| | | MED | | | | 0 (0%) | 5,125 (100%) | 0 (0%) | 615 (100%) | 0 (0%) | 4,510 (100%) |
| | Cephalochordata and Tunicata | ITA | 188 (0.24%) | 4 (2%) | 184 (98%) | 0 (0%) | 188 (100%) | 0 (0%) | 4 (100%) | 0 (0%) | 184 (100%) |
| | | EU | | | | 0 (0%) | 188 (100%) | 0 (0%) | 4 (100%) | 0 (0%) | 184 (100%) |
| | | GLO | | | | 0 (0%) | 188 (100%) | 0 (0%) | 4 (100%) | 0 (0%) | 184 (100%) |
| | | MED | | | | 0 (0%) | 188 (100%) | 0 (0%) | 4 (100%) | 0 (0%) | 184 (100%) |
| | Copepoda | ITA | 1,058 (1.38%) | 117 (11%) | 941 (89%) | 0 (0%) | 1,058 (100%) | 0 (0%) | 117 (100%) | 0 (0%) | 941 (100%) |
| | | EU | | | | 0 (0%) | 1,058 (100%) | 0 (0%) | 117 (100%) | 0 (0%) | 941 (100%) |
| | | GLO | | | | 2 (0%) | 1,056 (100%) | 2 (2%) | 115 (98%) | 0 (0%) | 941 (100%) |
| | | MED | | | | 0 (0%) | 1,058 (100%) | 0 (0%) | 117 (100%) | 0 (0%) | 941 (100%) |
| | Holothuroidea | ITA | 37 (0.05%) | - | 37 (100%) | 0 (0%) | 37 (100%) | - | - | 0 (0%) | 37 (100%) |
| | | EU | | | | 0 (0%) | 37 (100%) | - | - | 0 (0%) | 37 (100%) |
| | | GLO | | | | 6 (16%) | 31 (84%) | - | - | 6 (16%) | 31 (84%) |
| | | MED | | | | 0 (0%) | 37 (100%) | - | - | 0 (0%) | 37 (100%) |
| | Malacostraca | ITA | 1,616 (2.1%) | 390 (24%) | 1,226 (76%) | 0 (0%) | 1,616 (100%) | 0 (0%) | 390 (100%) | 0 (0%) | 1,226 (100%) |
| | | EU | | | | 0 (0%) | 1,616 (100%) | 0 (0%) | 390 (100%) | 0 (0%) | 1,226 (100%) |
| | | GLO | | | | 23 (1%) | 1,593 (99%) | 3 (1%) | 387 (99%) | 20 (2%) | 1,206 (98%) |

| | | | | | | | | | | | |
|---|---|---|---|---|---|---|---|---|---|---|---|
| | | MED | | | | 1 (0%) | 1,615 (100%) | 0 (0%) | 390 (100%) | 1 (0%) | 1,225 (100%) |
| | Other invertebrates | ITA | 8,518 (11.1%) | 1,086 (13%) | 7,432 (87%) | 0 (0%) | 8,518 (100%) | 0 (0%) | 1,086 (100%) | 0 (0%) | 7,432 (100%) |
| | | EU | | | | 0 (0%) | 8,518 (100%) | 0 (0%) | 1,086 (100%) | 0 (0%) | 7,432 (100%) |
| | | GLO | | | | 1 (0%) | 8,517 (99%) | 0 (0%) | 1,086 (100%) | 1 (0%) | 7,432 (100%) |
| | | MED | | | | 0 (0%) | 8,518 (100%) | 0 (0%) | 1,086 (100%) | 0 (0%) | 7,432 (100%) |
| **Vascular plants** | Gnetopsida | ITA | 9 (0.01%) | 2 (22%) | 7 (78%) | 4 (44%) | 5 (56%) | 0 (0%) | 2 (100%) | 4 (57%) | 3 (43%) |
| | | EU | | | | 2 (22%) | 7 (78%) | 0 (0%) | 2 (100%) | 2 (29%) | 5 (71%) |
| | | GLO | | | | 6 (67%) | 3 (33%) | 0 (0%) | 2 (100%) | 6 (86%) | 1 (14%) |
| | | MED | | | | 0 (0%) | 9 (100%) | 0 (0%) | 2 (100%) | 0 (0%) | 7 (100%) |
| | Liliopsida | ITA | 1,469 (1.91%) | 230 (16%) | 1,239 (84%) | 578 (39%) | 891 (61%) | 209 (91%) | 21 (9%) | 369 (30%) | 870 (70%) |
| | | EU | | | | 541 (37%) | 928 (63%) | 106 (46%) | 124 (54%) | 435 (35%) | 804 (65%) |
| | | GLO | | | | 444 (30%) | 1,025 (70%) | 87 (38%) | 143 (62%) | 357 (29%) | 882 (71%) |
| | | MED | | | | 366 (25%) | 1,103 (75%) | 96 (42%) | 134 (58%) | 270 (22%) | 969 (78%) |
| | Lycopodiopsida | ITA | 23 (0.03%) | 2 (9%) | 21 (91%) | 12 (52%) | 11 (48%) | 1 (50%) | 1 (50%) | 11 (52%) | 10 (48%) |
| | | EU | | | | 20 (87%) | 3 (13%) | 2 (100%) | 0 | 18 (86%) | 3 (14%) |
| | | GLO | | | | 8 (35%) | 15 (65%) | 2 (100%) | 0 | 6 (29%) | 15 (71%) |
| | | MED | | | | 7 (30%) | 16 (70%) | 2 (100%) | 0 | 5 (24%) | 16 (76%) |
| | Magnoliopsida | ITA | 7,361 (10%) | 1520 (21%) | 5,841 (79%) | 1,906 (26%) | 5,455 (74%) | 1,208 (79%) | 312 (21%) | 698 (12%) | 5,143 (88%) |
| | | EU | | | | 977 (13%) | 6,384 (87%) | 147 (10%) | 1,373 (90%) | 830 (14%) | 5,011 (86%) |
| | | GLO | | | | 688 (9%) | 6,673 (91%) | 144 (9%) | 1,376 (91%) | 544 (9%) | 5,297 (91%) |
| | | MED | | | | 207 (3%) | 7,154 (97%) | 63 (4%) | 1,457 (96%) | 144 (2%) | 5,697 (98%) |
| | Pinopsida | ITA | 25 (0.03%) | 2 (8%) | 23 (92%) | 4 (16%) | 21 (84%) | 2 (100%) | 0 | 2 (9%) | 21 (91%) |
| | | EU | | | | 23 (92%) | 2 (8%) | 2 (100%) | 0 | 21 (91%) | 2 (9%) |
| | | GLO | | | | 23 (92%) | 2 (8%) | 2 (100%) | 0 | 21 (91%) | 2 (9%) |

| | | | | | | | | | | | |
|---|---|---|---|---|---|---|---|---|---|---|---|
| | | MED | | | | 1 (4%) | 24 (96%) | 1 (50%) | 1 (50%) | 0 (0%) | 23 (100%) |
| | Polypodiopsida | ITA | 115 (0.15%) | 1 (1%) | 114 (99%) | 20 (17%) | 95 (83%) | 1 (100%) | 0 | 19 (17%) | 95 (83%) |
| | | EU | | | | 100 (87%) | 15 (13%) | 0 (0%) | 1 (100%) | 100 (88%) | 14 (12%) |
| | | GLO | | | | 20 (17%) | 95 (83%) | 0 (0%) | 1 (100%) | 20 (18%) | 94 (82%) |
| | | MED | | | | 8 (7%) | 107 (93%) | 0 (0%) | 1 (100%) | 8 (7%) | 106 (93%) |
| **Vertebrates** | Actinopterygii | ITA | 443 (1%) | 22 (5%) | 421 (95%) | 434 (98%) | 9 (2%) | 22 (100%) | 0 | 412 (98%) | 9 (2%) |
| | | EU | | | | 428 (97%) | 15 (3%) | 21 (95%) | 1 (5%) | 407 (97%) | 14 (3%) |
| | | GLO | | | | 387 (88%) | 56 (13%) | 21 (95%) | 1 (5%) | 366 (87%) | 55 (13%) |
| | | MED | | | | 362 (82%) | 81 (18%) | 9 (41%) | 13 (59%) | 353 (84%) | 68 (16%) |
| | Amphibia | ITA | 49 (0.06%) | 20 (41%) | 29 (59%) | 47 (96%) | 2 (4%) | 20 (100%) | 0 | 27 (93%) | 2 (7%) |
| | | EU | | | | 48 (98%) | 1 (2%) | 20 (100%) | 0 | 28 (97%) | 1 (3%) |
| | | GLO | | | | 45 (92%) | 4 (8%) | 19 (95%) | 1 (5%) | 26 (90%) | 3 (10%) |
| | | MED | | | | 11 (22%) | 38 (78%) | 9 (45%) | 11 (55%) | 2 (7%) | 27 (93%) |
| | Aves | ITA | 343 (0.45%) | - | 343 (100%) | 310 (90%) | 33 (10%) | - | - | 310 (90%) | 33 (10%) |
| | | EU | | | | 340 (99%) | 3 (1%) | - | - | 340 (99%) | 3 (1%) |
| | | GLO | | | | 339 (99%) | 4 (1%) | - | - | 339 (99%) | 4 (1%) |
| | | MED | | | | 42 (12%) | 301 (88%) | - | - | 42 (12%) | 301 (88%) |
| | Chondrichthyes | ITA | 69 (0.1%) | - | 69 (100%) | 66 (96%) | 3 (4%) | - | - | 66 (96%) | 3 (4%) |
| | | EU | | | | 64 (93%) | 5 (7%) | - | - | 64 (93%) | 5 (7%) |
| | | GLO | | | | 65 (94%) | 4 (6%) | - | - | 65 (94%) | 4 (6%) |
| | | MED | | | | 58 (84%) | 11 (16%) | - | - | 58 (84%) | 11 (16%) |
| | Agnatha | ITA | 5 (0.01%) | 1 (20%) | 4 (80%) | 4 (80%) | 1 (20%) | 1 (100%) | 0 | 3 (75%) | 1 (25%) |
| | | EU | | | | 5 (100%) | 0 (0%) | 1 (100%) | 0 | 4 (100%) | 0 (0%) |
| | | GLO | | | | 5 (100%) | 0 (0%) | 1 (100%) | 0 | 4 (100%) | 0 (0%) |
| | | MED | | | | 1 (20%) | 4 (80%) | 0 (0%) | 1 (100%) | 1 (25%) | 3 (75%) |
| | Mammalia | ITA | 165 (0.2%) | 42 (25%) | 123 (75%) | 152 (92%) | 13 (8%) | 42 (100%) | 0 | 110 (89%) | 13 (11%) |

| | | | | | | | | | | | |
|---|---|---|---|---|---|---|---|---|---|---|---|
| | | EU | | | | 159 (96%) | 6 (4%) | 40 (95%) | 2 (5%) | 119 (97%) | 2 (5%) |
| | | GLO | | | | 152 (92%) | 13 (8%) | 39 (93%) | 3 (7%) | 113 (92%) | 10 (8%) |
| | | MED | | | | 108 (65%) | 57 (35%) | 30 (71%) | 12 (29%) | 78 (63%) | 45 (37%) |
| | Reptilia | ITA | 64 (0.08%) | 13 (20%) | 51 (80%) | 56 (88%) | 8 (14%) | 12 (92%) | 1 (8%) | 44 (86%) | 7 (14%) |
| | | EU | | | | 56 (88%) | 8 (14%) | 12 (92%) | 1 (8%) | 44 (86%) | 7 (14%) |
| | | GLO | | | | 57 (89%) | 8 (12%) | 12 (92%) | 1 (8%) | 45 (88%) | 6 (12%) |
| | | MED | | | | 7 (11%) | 57 (89%) | 4 (31%) | 9 (69%) | 3 (6%) | 48 (94%) |
| **Major taxa** | **Taxa groups** | **Scope** | **Totals per major group** | **Total endemics** | **Total non endemics** | **Total assessed** | **Total non-assessed** | **Assessed endemics** | **Non assessed endemics** | **Assessed non-endemics** | **Non assessed non endemics** |
| ***Totals*** | | | 76,845 | 8,389 (11%) | 68,456 (89%) | 7,349 (10%) | 69,496 (90%) | 1,700 (20%) | 6,689 (80%) | 5,649 (8%) | 62,807 (92%) |
| | | | | | | 7,273 (10%) | 69,572 (91%) | 832 (10%) | 7,557 (90%) | 6,441 (9%) | 62,015 (91%) |
| | | | | | | 3,820 (5%) | 73,025 (95%) | 790 (9%) | 7,599 (91%) | 3,030 (4%) | 65,426 (96%) |
| | | | | | | 1,800 (2%) | 75,045 (98%) | 418 (5%) | 7,971 (95%) | 1,382 (2%) | 67,074 (98%) |
