## Supplementary Table 3 for "How well is Italian biodiversity represented in red lists and conservation legislation? Taxonomic biases, coverage gaps and the assessment-to-legislation bottleneck"

**Supplementary table 3:** Summary of counts per major taxa included (Yes) or not included (No) in any policy for each IUCN category.

| | | | Yes | No | Total |
|---|---|---|---|---|---|
| **Insects** | ***Coleoptera*** | | **11** | **1,738** | **1,749** |
| | | **CR** | | 70 | 70 |
| | | **DD** | | 189 | 189 |
| | | **EN** | 5 | 102 | 107 |
| | | **LC** | 2 | 881 | 883 |
| | | **NT** | 1 | 300 | 301 |
| | | **RE** | | 2 | 2 |
| | | **VU** | 3 | 194 | 197 |
| | ***Hymenoptera*** | | | **124** | **124** |
| | | **CR** | | 1 | 1 |
| | | **DD** | | 94 | 94 |
| | | **EN** | | 8 | 8 |
| | | **NT** | | 12 | 12 |
| | | **PE** | | 6 | 6 |
| | | **VU** | | 3 | 3 |
| | ***Lepidoptera*** | | **20** | **248** | **268** |
| | | **CR** | 1 | | 1 |
| | | **DD** | | 1 | 1 |
| | | **EN** | 3 | 5 | 8 |
| | | **LC** | 10 | 225 | 235 |
| | | **NT** | 2 | 13 | 15 |
| | | **RE** | 1 | | 1 |
| | | **VU** | 3 | 4 | 7 |
| | ***Odonata*** | | **9** | **74** | **83** |
| | | **CR** | 1 | 1 | 2 |
| | | **DD** | 1 | 2 | 3 |
| | | **EN** | 1 | 3 | 4 |
| | | **LC** | 1 | 60 | 61 |
| | | **NT** | 5 | 4 | 9 |
| | | **VU** | | 4 | 4 |
| | **Total** | | **40** | **2,184** | **2,224** |
| **Lichens** | ***Ascolichen*** | | **29** | **125** | **154** |
| | | **CR** | | 18 | 18 |
| | | **DD** | 9 | 4 | 13 |
| | | **EN** | | 15 | 15 |
| | | **LC** | 2 | 5 | 7 |
| | | **NT** | 8 | 17 | 25 |
| | | **RE** | | 29 | 29 |
| | | **VU** | 10 | 37 | 47 |
| | **Total** | | **29** | **125** | **154** |
| **Macrofungi** | ***Ascomycota*** | | | **2** | **2** |
| | | **CR** | | 1 | 1 |
| | | **VU** | | 1 | 1 |
| | ***Basidiomycota*** | | | **19** | **19** |
| | | **CR** | | 7 | 7 |

| | | | | | |
|---|---|---|---|---|---|
| | | **DD** | | 2 | 2 |
| | | **EN** | | 5 | 5 |
| | | **NT** | | 3 | 3 |
| | | **VU** | | 2 | 2 |
| | **Total** | | | **21** | **21** |
| **Non-vascular plants** | ***Andreaeopsida*** | | | **10** | **10** |
| | | **DD** | | 3 | 3 |
| | | **LC** | | 3 | 3 |
| | | **NT** | | 1 | 1 |
| | | **VU** | | 3 | 3 |
| | ***Anthocerotopsida*** | | **1** | **5** | **6** |
| | | **CR** | | 1 | 1 |
| | | **LC** | | 3 | 3 |
| | | **NT** | | 1 | 1 |
| | | **PE** | 1 | | 1 |
| | ***Bryopsida*** | | **9** | **880** | **889** |
| | | **CR** | 1 | 31 | 32 |
| | | **DD** | 1 | 93 | 94 |
| | | **EN** | 1 | 102 | 103 |
| | | **LC** | 3 | 493 | 496 |
| | | **NT** | 2 | 50 | 52 |
| | | **PE** | 1 | 14 | 15 |
| | | **RE** | | 1 | 1 |
| | | **VU** | | 96 | 96 |
| | ***Marchantiophyta*** | | **6** | **291** | **297** |
| | | **CR** | 2 | 12 | 14 |
| | | **DD** | | 32 | 32 |
| | | **EN** | 3 | 35 | 38 |
| | | **LC** | | 148 | 148 |
| | | **NT** | 1 | 27 | 28 |
| | | **PE** | | 2 | 2 |
| | | **RE** | | 4 | 4 |
| | | **VU** | | 31 | 31 |
| | ***Polytrichopsida*** | | | **21** | **21** |
| | | **DD** | | 3 | 3 |
| | | **EN** | | 1 | 1 |
| | | **LC** | | 15 | 15 |
| | | **NT** | | 1 | 1 |
| | | **VU** | | 1 | 1 |
| | ***Sphagnopsida*** | | **34** | | **34** |
| | | **CR** | 4 | | 4 |
| | | **DD** | 1 | | 1 |
| | | **EN** | 2 | | 2 |
| | | **LC** | 21 | | 21 |
| | | **NT** | 2 | | 2 |
| | | **VU** | 4 | | 4 |
| | **Total** | | **50** | **1,207** | **1,257** |
| **Other invertebrates** | ***Anthozoa*** | | **33** | **67** | **100** |
| | | **CR** | 3 | 1 | 4 |

| | | | | | |
|---|---|---|---|---|---|
| | | **DD** | 20 | 43 | 63 |
| | | **EN** | 1 | | 1 |
| | | **LC** | 5 | 21 | 26 |
| | | **NT** | 2 | | 2 |
| | | **VU** | 2 | 2 | 4 |
| **Vascular plants** | ***Gnetopsida*** | | | **4** | **4** |
| | | **DD** | | 3 | 3 |
| | | **EN** | | 1 | 1 |
| | ***Liliopsida*** | | **176** | **402** | **578** |
| | | **CR** | 5 | 16 | 21 |
| | | **DD** | 15 | 43 | 58 |
| | | **EN** | 21 | 44 | 65 |
| | | **LC** | 102 | 186 | 288 |
| | | **NT** | 22 | 76 | 98 |
| | | **PE** | 4 | 5 | 9 |
| | | **PEW** | | 1 | 1 |
| | | **VU** | 7 | 31 | 38 |
| | ***Lycopodiopsida*** | | **3** | **9** | **12** |
| | | **CR** | 1 | | 1 |
| | | **DD** | 2 | 5 | 7 |
| | | **LC** | | 3 | 3 |
| | | **NT** | | 1 | 1 |
| | ***Magnoliopsida*** | | **102** | **1,804** | **1,906** |
| | | **CR** | 13 | 113 | 126 |
| | | **DD** | 3 | 352 | 355 |
| | | **EN** | 25 | 179 | 204 |
| | | **EW** | 1 | 2 | 3 |
| | | **EX** | | 8 | 8 |
| | | **LC** | 25 | 729 | 754 |
| | | **NT** | 22 | 276 | 298 |
| | | **PE** | 4 | 27 | 31 |
| | | **RE** | 1 | | 1 |
| | | **VU** | 8 | 118 | 126 |
| | ***Pinopsida*** | | **1** | **3** | **4** |
| | | **CR** | 1 | | 1 |
| | | **LC** | | 2 | 2 |
| | | **NT** | | 1 | 1 |
| | ***Polypodiopsida*** | | **11** | **9** | **20** |
| | | **CR** | 3 | 1 | 4 |
| | | **DD** | | 1 | 1 |
| | | **EN** | 4 | 2 | 6 |
| | | **LC** | 2 | 1 | 3 |
| | | **NT** | | 2 | 2 |
| | | **PE** | | 1 | 1 |
| | | **VU** | 2 | 1 | 3 |
| | **Total** | | **293** | **2,231** | **2,524** |
| **Vertebrates** | ***Actinopterygii*** | | **43** | **391** | **432** |
| | | **CR** | 8 | 5 | 13 |
| | | **DD** | 4 | 48 | 52 |

| | | | | | |
|---|---|---|---|---|---|
| | | **EN** | 8 | 4 | 12 |
| | | **LC** | 9 | 322 | 331 |
| | | **NT** | 7 | 6 | 13 |
| | | **RE** | 2 | | 2 |
| | | **VU** | 5 | 6 | 11 |
| | ***Agnatha*** | | **4** | | **4** |
| | | **CR** | 2 | | 2 |
| | | **VU** | 2 | | 2 |
| | ***Amphibia*** | | **46** | **1** | **47** |
| | | **CR** | 1 | | 1 |
| | | **EN** | 7 | | 7 |
| | | **LC** | 23 | | 23 |
| | | **NT** | 8 | | 8 |
| | | **VU** | 7 | 1 | 8 |
| | ***Aves*** | | **296** | **14** | **310** |
| | | **CR** | 11 | | 11 |
| | | **DD** | 15 | | 15 |
| | | **EN** | 26 | | 26 |
| | | **LC** | 163 | 11 | 174 |
| | | **NT** | 40 | 2 | 42 |
| | | **RE** | 4 | | 4 |
| | | **VU** | 37 | 1 | 38 |
| | ***Chondrichthyes*** | | **39** | **27** | **66** |
| | | **CR** | 10 | | 10 |
| | | **DD** | 20 | 10 | 30 |
| | | **EN** | 5 | | 5 |
| | | **LC** | 2 | 13 | 15 |
| | | **NT** | 1 | 3 | 4 |
| | | **VU** | 1 | 1 | 2 |
| | ***Mammalia*** | | **110** | **42** | **152** |
| | | **CR** | 3 | | 3 |
| | | **DD** | 11 | 5 | 16 |
| | | **EN** | 10 | 1 | 11 |
| | | **LC** | 59 | 33 | 92 |
| | | **NT** | 15 | 3 | 18 |
| | | **VU** | 12 | 1 | 12 |
| | ***Reptilia*** | | **51** | **5** | **56** |
| | | **CR** | | 1 | 1 |
| | | **DD** | 1 | 1 | 1 |
| | | **EN** | 8 | | 8 |
| | | **LC** | 36 | 3 | 39 |
| | | **NT** | 4 | 1 | 5 |
| | | **VU** | 2 | | 2 |
| | **Total** | | **590** | **479** | **1,069** |
| **Total** | | | 1,035 | 6,314 | 7,349 |

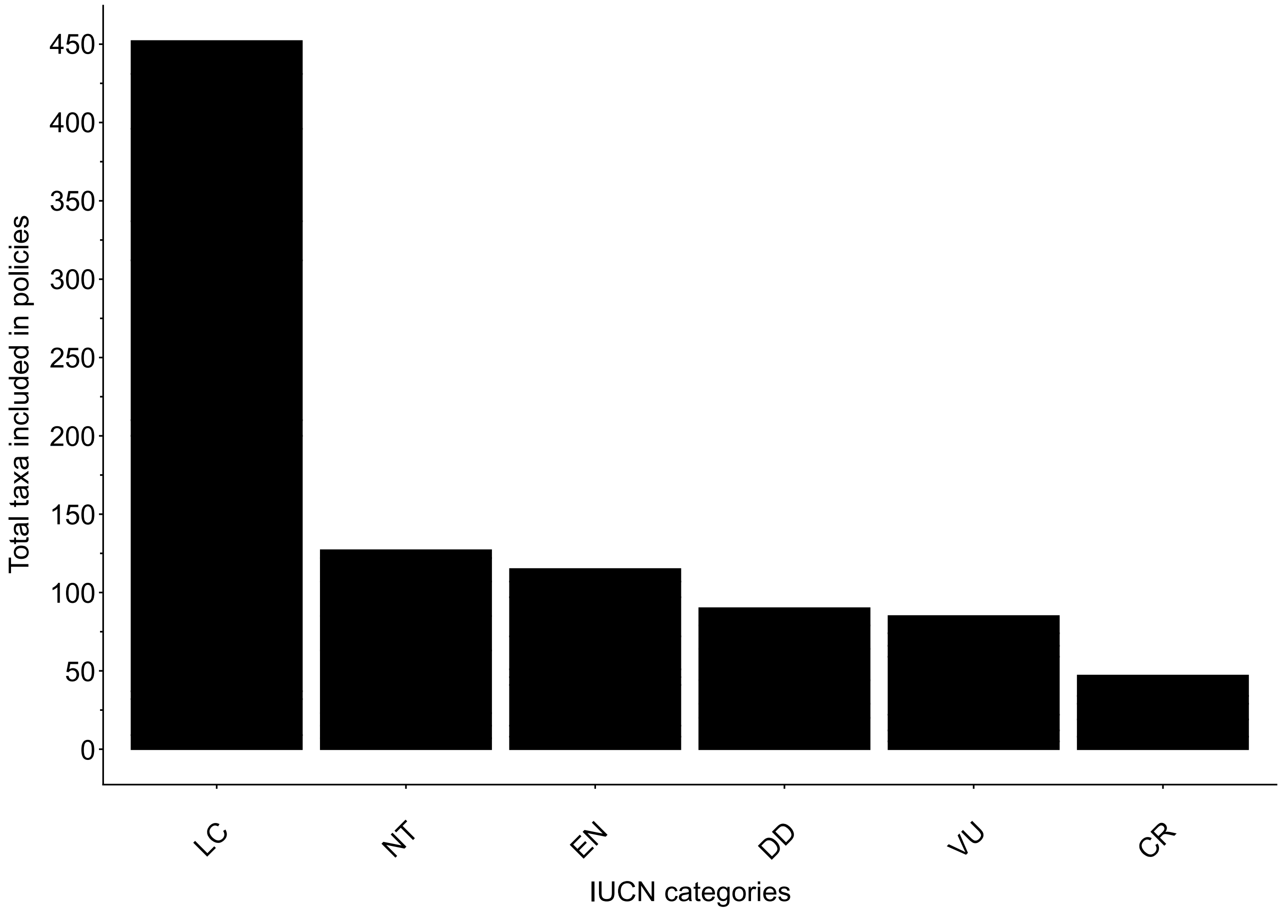


**Supplementary figure 3:** Total counts of taxa per IUCN category in Italian Red Lists included in any policy, with bars ordered according to highest count per category (LC = Least Concern, NT = Near threatened, DD = Data Deficient, EN = Endangered, VU = Vulnerable, CR = Critically Endangered).

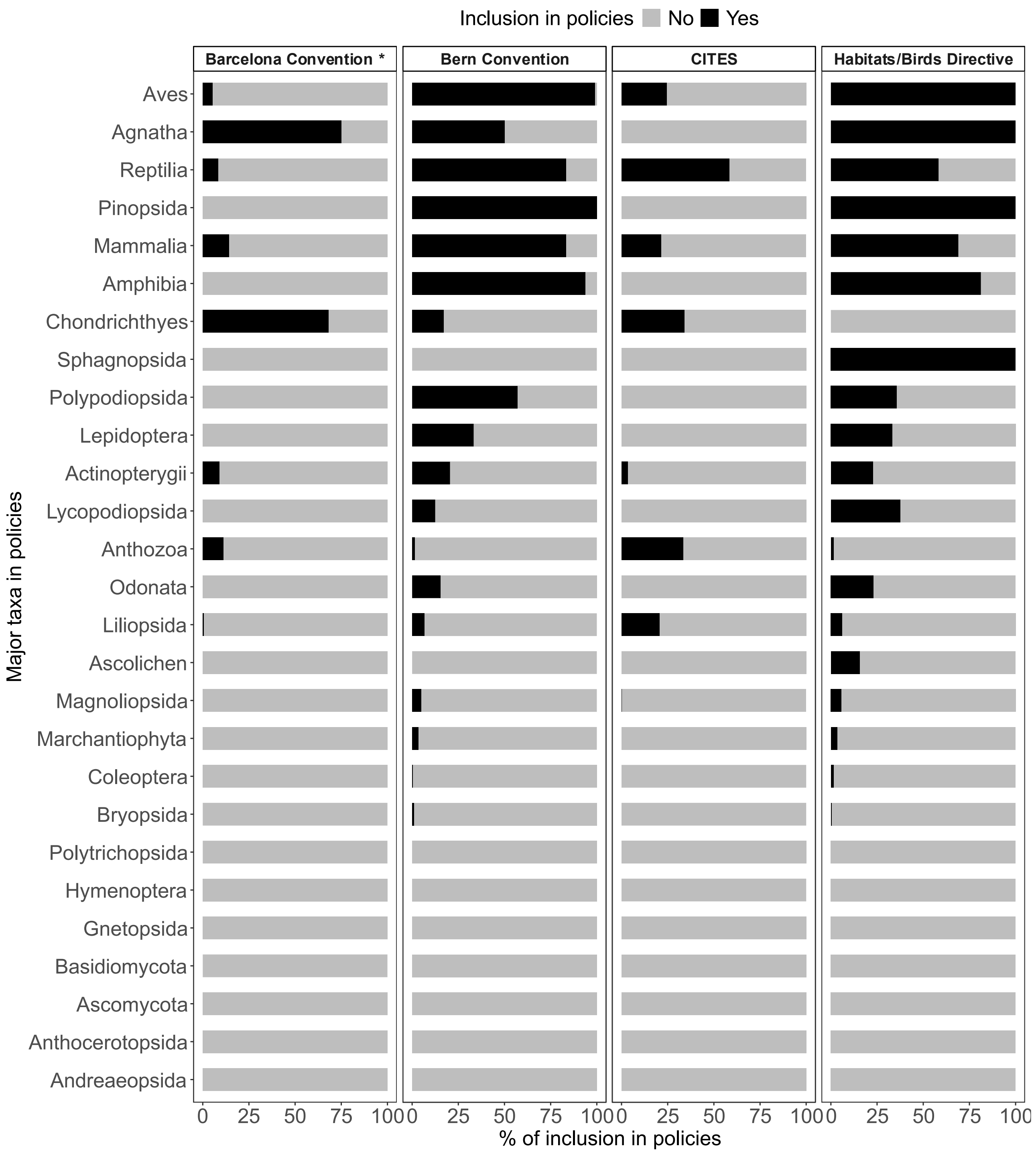


**Supplementary figure 4:** percentages of threatened taxa included vs excluded in the appendixes of legislation reviewed in the manuscript.

*the Barcelona Convention focuses only on marine taxa

**Supplementary table 4:** Results of Fisher's exact test for threatened taxa (a) and endemic taxa (b). The table showed p-values in the first row and the summary results of test including odds-ratio (OR), low and upper confidence intervals and counts.

a)

| *Taxa in policies* | *p* | *OR* | *OR_CI_L* | *OR_CI_H* | *Non_threat_not in policies* | *Non_threat_in policies* | *Threat_not in policies* | *Threat_in policies* |
|---|---|---|---|---|---|---|---|---|
| *Actinopterygii* | <0.001 | 29.92 | 9.52 | 99.07 | 328 | 13 | 9 | 11 |
| *Chondrichthyes* | <0.001 | 68.49 | 6.79 | 3676.13 | 16 | 3 | 1 | 16 |
| *Lepidoptera* | <0.001 | 17.32 | 4.09 | 72.79 | 229 | 11 | 7 | 6 |
| *Coleoptera* | <0.01 | 8.41 | 1.91 | 50.72 | 1120 | 3 | 310 | 7 |
| *Ascolichen* | <0.01 | 0.23 | 0.07 | 0.68 | 22 | 10 | 99 | 10 |
| *Marchantiophyta* | 0.02 | 10.32 | 1.13 | 494.12 | 175 | 1 | 84 | 5 |
| *Mammalia* | 0.02 | 7.89 | 1.12 | 345.87 | 22 | 55 | 1 | 20 |
| *Anthozoa* | 0.04 | 6.92 | 0.91 | 89.42 | 15 | 6 | 2 | 6 |
| *Aves* | 0.12 | 4.98 | 0.72 | 214.77 | 13 | 203 | 1 | 78 |
| *Magnoliopsida* | 0.14 | 1.63 | 0.84 | 3.14 | 350 | 24 | 188 | 21 |
| *Odonata* | 0.21 | 3.14 | 0.26 | 23.62 | 64 | 5 | 8 | 2 |
| *Liliopsida* | 0.59 | 0.83 | 0.46 | 1.46 | 162 | 65 | 75 | 25 |
| *Polypodiopsida* | 0.61 | 2.28 | 0.19 | 36.86 | 3 | 2 | 5 | 8 |
| *Bryopsida* | 0.71 | 1.33 | 0.21 | 6.92 | 543 | 5 | 244 | 3 |

b)

| *Taxa in policies* | *p* | *OR* | *OR_CI_L* | *OR_CI_H* | *End_policies* | *End_not in policies* | *Non-end_policies* | *Non-end_not in policies* |
|---|---|---|---|---|---|---|---|---|
| *Actinopterygii* | <0.001 | 28.41 | 10 | 89 | 15 | 7 | 29 | 392 |
| *Liliopsida* | <0.001 | 3 | 2.16 | 4.14 | 80 | 150 | 187 | 1,052 |
| *Magnoliopsida* | <0.001 | 3.2 | 2.18 | 4.64 | 55 | 1465 | 68 | 5,773 |
| *Anthozoa* | 0.01 | 0.2 | 0.04 | 0.73 | 3 | 29 | 32 | 62 |
| *Mammalia* | 0.03 | 0.4 | 0.17 | 0.9 | 25 | 17 | 97 | 26 |
| *Lepidoptera* | 0.04 | 4.14 | 0.79 | 14 | 3 | 157 | 23 | 4,985 |
| *Gastropoda* | 0.09 | 0.18 | 0.004 | 1.18 | 1 | 537 | 16 | 1,563 |
| *Lycopodiopsida* | 0.25 | 7.96 | 0.08 | 776.6 | 1 | 1 | 2 | 19 |
| *Reptilia* | 0.37 | 0.45 | 0.08 | 3.26 | 10 | 3 | 45 | 6 |
| *Malacostraca* | 0.47 | 0.35 | 0.008 | 2.52 | 1 | 389 | 9 | 1,217 |
| *Coleoptera* | 0.75 | 0.58 | 0.06 | 2.51 | 2 | 2606 | 14 | 10,526 |
| *Porifera* | 0.77 | 0.76 | 0.16 | 3.03 | 4 | 207 | 7 | 275 |
